# Shugoshin promotes efficient activation of spindle assembly checkpoint and timely spindle disassembly

**DOI:** 10.1101/2020.09.04.282871

**Authors:** Aakanksha Sane, Shreyas Sridhar, Kaustuv Sanyal, Santanu K Ghosh

## Abstract

Shugoshin proteins are evolutionary conserved across eukaryotes with some species-specific cellular functions ensuring the fidelity of chromosome segregation. Shugoshin being present at various subcellular locales, acts as an adaptor to mediate various protein-protein interactions in a spatio-temporal manner. Here, we characterize shugoshin (Sgo1) in the human fungal pathogen, *Candida albicans*. Interestingly, we discover a novel *in vivo* localization of Sgo1 along the length of the mitotic spindle. Further, Sgo1 performs a hitherto unknown function of facilitating timely disassembly of spindle in this organism. We observe that Sgo1 retains its centromeric localization and performs its conserved functions that include regulating the centromeric condensin localization, chromosome passenger complex (CPC) maintenance and sister chromatid biorientation. We identify novel roles of Sgo1 as a spindle assembly checkpoint (SAC) component with functions in maintaining the SAC proteins, Mad2 and Bub1, at the kinetochores, in response to faulty kinetochore-microtubule attachments. These findings provide an excellent evidence of the functional rewiring of shugoshin in maintaining genomic stability.

## Introduction

The bipolar attachment of sister chromatids to opposite spindle poles is mediated by dynamic interactions between kinetochores and microtubules during mitosis. Such biorientation keeps the sister chromatids under tension. by counter-balancing opposing pulling forces. The force towards the spindle poles generated by microtubule depolymerization is opposed by the cohesion formed between the sister chromatids (reviewed in Cleveland et al 2003, Maiato et al 2004, Tanaka et al 2005). Any failure in the kinetochore-microtubule (KT-MT) attachment and a lack of tension between the sisters is sensed by a network of proteins including the spindle assembly checkpoint (SAC) which then, in response, elicits a cell cycle arrest at the metaphase stage (reviewed in Vleugel et al 2012, Musacchio 2015). Although, the mechanism of arrest through inactivation of Cdc20, a co-activator of the anaphase promoting complex (APC) is well elucidated, the molecular details of how the proper attachment and tension are sensed are not fully understood.

*Drosophila* Mei-S332 was first discovered and predicted to be a protector of centromeric cohesion during meiosis I (Kerrebrock et al 1992, Kerrebrock et al 1995). Later, its distant relative shugoshin was discovered (Kitajima et al 2004) as the protector of centromeric cohesion at metaphase I to ensure non-disjunction of the sister chromatids during meiosis I which is crucial for reductional division. This function has been found to be conserved across sexually reproducing organisms (Kerrebrock et al 1992, Kitajima et al 2004, Katis et al 2004, Marston et al 2004, Rabitsch et al 2004, Hamant et al 2005, Wang et al 2011, Cromer et al 2013, Llano et al 2008). The shugoshin family of proteins, has also been implicated in sensing the tension between the sister chromatids in various organisms (Indjeian et al 2005, Kawashima et al 2007, Huang et al 2007). Sgo2, a paralog of shugoshin (Sgo1), is present in several organisms including *Schizosaccharomyces pombe* (Kitajima et al 2004). To protect centromeric cohesion, shugoshin acts as an adapter to recruit protein phosphatase 2A (PP2A) at the centromeres and dephosphorylates meiotic cohesin Rec8 to prevent it from being cleaved by separase (Riedel et al 2006, Kitajima et al 2006, Xu et al 2009). Likewise, shugoshin has also been shown to protect centromeric cohesin from removal by ‘prophase pathway’ during mitosis (McGuinness et al 2005, Rivera and Losada 2009). Since its discovery, the adapter function of shugoshin has been shown to recruit various proteins at and around the centromeres (pericentromeres) to cater different centromere related functions across eukaryotes, often in a species-specific manner. Recently, in the budding yeast *Saccharomyces cerevisiae*, ScSgo1 has been shown to promote mono-orientation of the sister chromatids during meiosis I by facilitating monopolin localization at the centromeres (Mehta et al 2018).

In mitosis, the function of shugoshin as a sensor of tension between the sister chromatids is believed to be driven by the enrichment of this protein at the centromeres that are not under tension (Kiburz et al 2005, Nerusheva et al 2014). Centromere-localized shugoshin then in turn promotes correction of erroneous KT-MT attachment and biorientation of the sister chromatids on the microtubule spindle through recruitment of Chromosomal Passenger Complex (CPC), PP2A and condensin at the centromeres (Xu et al 2009, Eshleman et al 2014, Verzijlbergen et al 2014, Peplowska et al 2014, Nerusheva et al 2014). Besides sensing the lack of tension, whether shugoshin relays the signal downstream by activating SAC remains unclear. In budding yeast, Sgo1 is dispensable for the growth and also for mounting cell cycle arrest in response to lack of tension. However, a significant drop in cell viability observed in this organism for *sgo1-100* mutant cells during recovery from a spindle poison is believed to be due to the failure in *de novo* establishment of sister chromatid biorientation (Indjeian et al 2005). On the other hand, shugoshin has been shown to interact *in vivo* with the SAC component Mad2 in humans, mice and frogs. Notably, although mouse/human Sgo2 and frog Sgo1 exhibit a Mad1/Cdc20-like interaction with Mad2, this interaction is dispensable for SAC activation in mitosis. Rather this interaction has been implicated in meiosis for a possible SAC signaling and cell cycle arrest during meiosis II (Orth et al 2011, Hellmuth et al 2020). However, in support of an indirect promotion of SAC activation in mitosis, it has been demonstrated in budding yeast that ScSgo1 is involved in preventing SAC silencing in cells with unattached chromosomes (Jin and Wang 2013). Besides its roles at the KT-MT interface, other functions of shugoshin include promotion of homolog pairing by protecting centromeric synaptonemal complex (deAlmeida et al 2019). Interestingly, the adapter function of shugoshin is not only restricted at centromere/pericentromere but has been documented also at the chromosomal arm regions and even at the subtelomeric regions. Recently, Sgo1 has been implicated in chromosomal arm condensation in *S. cerevisiae* indicating possible localization of ScSgo1 along the arms (Kruitwagen et al 2018). In *C. elegans*, shugoshin has been shown to be essential for meiotic prophase checkpoint perhaps through its interaction with cohesin along the chromosomes (Bohr et al 2018). In *S. pombe*, shugoshin relocalizes from centromeres to the vicinity of telomeres possibly to regulate subtelomeric gene repression and late firing of replication origins (Tashiro et al 2016). Remarkably, in HeLa cells, a spliced variant of shugoshin (sSgo1) has also been observed at the spindle poles and mitotic spindles, where its function is believed to protect precocious sister centrioles disengagements (Wang et al 2006, Wang et al 2008, Mohr et al 2015).

*Candida albicans*, is the most prevalent fungal pathogen in humans (reviewed in Legrand et al 2019). This resides as a commensal organism in gastrointestinal and genitourinary tracts as a part of natural microflora but also as an opportunistic organism it infects mucous membrane, skin and nails and causes several non-lethal diseases collectively called as candidiasis (Sternberg 1994, Chin et al 2016). In immunocompromised patients, *C. albicans* can enter the bloodstream to cause systemic infection, where mortality rate can go up to 50% (Gudlaugsson et al 2003, Eggimann et al 2003). To strive with extreme environmental stresses within the host, *C. albicans* exhibits genome plasticity generating an endurable aneuploidy which is believed to also confer resistance to anti-fungal drugs (Reviewed in Sanyal 2012; Legrand et al 2019). Since the functions of shugoshin in accurate chromosome segregation appear to be conserved across the organisms with some species specific modulation, we wished to examine its roles in *C. albicans* for the first time to investigate whether and how aneuploidy can be generated due to alteration of its function. We observed that unlike several organisms but like *S. cerevisiae*, *C. albicans* harbors a single ortholog of shugoshin protein. While several functions including its non-essentiality under normal conditions were found to be conserved in *C. albicans*, important rewiring of functions in the form of SAC activation and spindle disassembly were also observed. Our results suggest that the loss of these functions under stress conditions, possibly within human host, has a potential to generate aneuploidy. Thus, this study uncovers yet other novel functions of shugoshin showing evolution of this protein in a human pathogen.

## Results

### Identification of Sgo1 in *C. albicans*

*SGO* genes although, are highly conserved among eukaryotes, exhibit a very low overall homology at the amino acid sequence (Kitajima et al 2004, reviewed in Marston 2015). The shugoshin family of proteins exhibit a conserved coiled coil domain at the N-terminus and a conserved basic region called ‘SGO’ motif near the C-terminus. We retrieved amino acid sequences of *ORF 19.3550* annotated as *SGO1* from <u>C</u>andida <u>G</u>enome <u>D</u>atabase (CGD; www.candidagenome.org) and of the *SGO* genes from various other eukaryotes. Using conventional CLUSTALW, GBLOCKS (Castresana 2000) and BLAST programs, we independently identified the conserved N-terminal domain (residues 26-103) and the SGO motif (residues 326-364) in *ORF 19.3550* to assign the same as Sgo1 in *C. albicans* (CaSgo1) (Suppl. Fig. S1A). The alignment of Sgo proteins from various organisms depicts the relative positions of these conserved regions (Suppl. Fig. S1B). We constructed a phylogenetic tree of shugoshin, using CLUSTAL Omega and Interactive Tree of Life (iTOL) (Ciccarelli et al 2006, Letunic et al 2006), to determine the evolutionary position of CaSgo1 with respect to other ascomycetes species (Suppl. Fig. S1C). The tree suggests that CaSgo1 is evolutionarily closer to ScSgo1 as there is a recent common ancestor which becomes evident when we compared the sequences of CaSgo1, ScSgo1 and SpSgo1 (Suppl. Fig. S1D). In several species, Sgo proteins are believed to be degraded by ubiquitination (Salic et al 2004, Karamysheva et al 2009). Ubiquitin-mediated degradation of proteins requires the presence of specific destruction motifs like D box and KEN box in the target proteins that are recognized by the ubiquitin ligases for degradation (Glotzer et al 1991, Pfleger and Kirschner 2000). *Homo sapiens* Sgo1 (HsSgo1) contains destruction motifs KEN box and R-L-N D box for degradation (Karamysheva et al 2009). ScSgo1 displays a modified KEN box as ‘NKSEN’ sequence (Eshleman et al 2014). Although in CaSgo1 we failed to identify a KEN box or the amino acid sequence NKSEN but it possesses a R-L-N D box motif (residues 402-410) composed of the sequence R-X-X-L-X-X-X-X-N (Fig. 1A) similar to that found in HsSgo1 which is one of the substrates for APC/C^Cdh1^ mediated degradation during mitotic exit (Karamysheva et al 2009). In addition, we detected a spindle-associated motif in CaSgo1 (residues 258-358), partly overlapping with the SGO motif, predicted by MOTIF (Kanehisa et al 2002) with an E-value of 0.68 (Fig. 1A). Since, many organisms including the unicellular fission yeast *S. pombe* and multicellular eukaryotes like mammals, contain two functionally distinct paralogs of shugoshin, namely *SGO1* and *SGO2,* we searched for an *SGO2* ortholog in CGD but failed to identify one. We thus report that like *S. cerevisiae*, *C. albicans* has a single shugoshin homolog *SGO1.*

**Figure 1.**
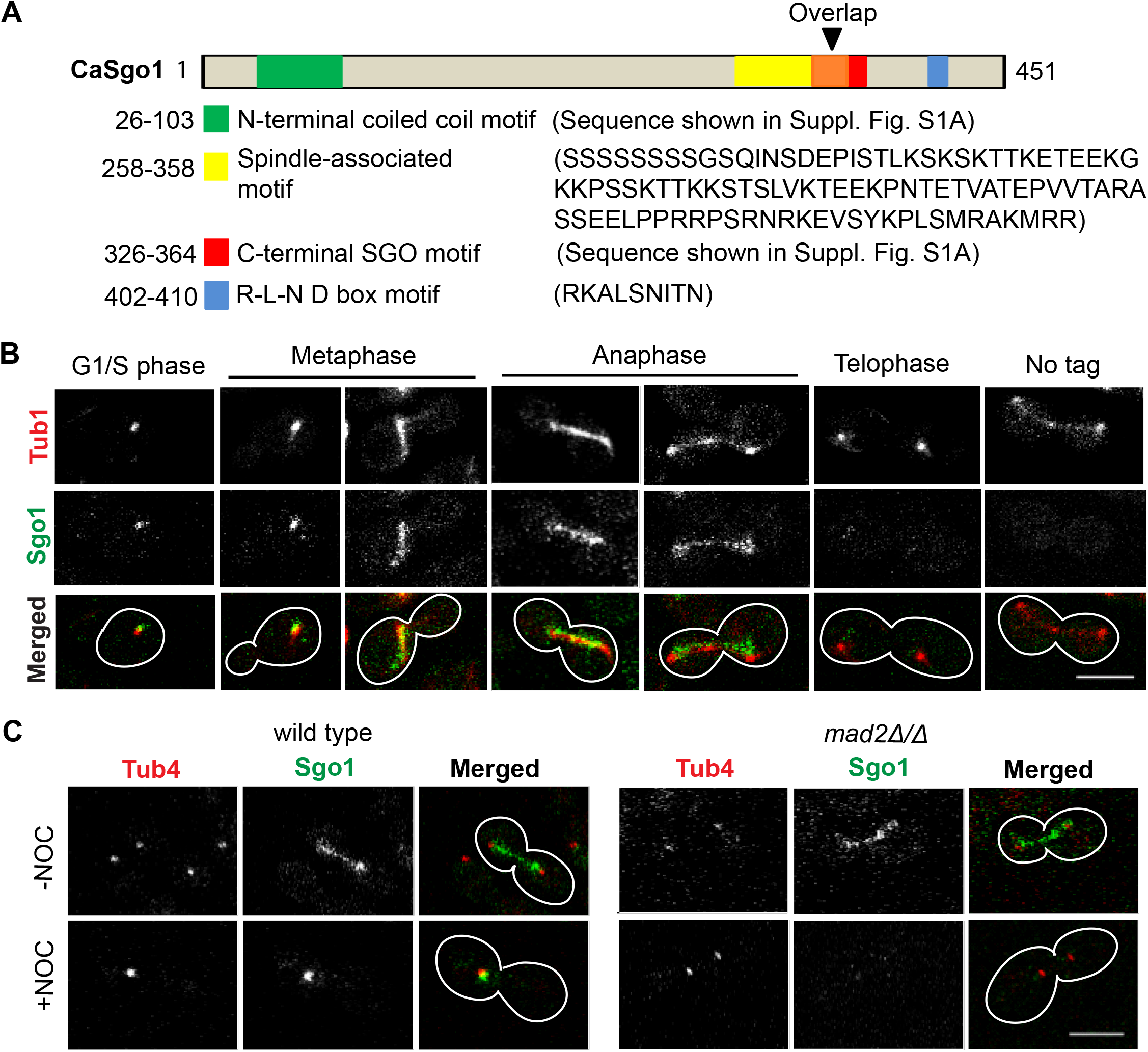
Cell cycle dependent localization of Sgo1 at the kinetochore and along the mitotic spindle. (A) In addition to N-terminal (green box) and C-terminal (red box) conserved motifs present in the shugoshin family of proteins, CaSgo1 possesses a spindle-associated motif (yellow box) predicted using the MOTIF sequence analysis tool and a conserved R-L-N D box motif (blue box). ‘Overlap’ (orange box, black arrowhead) is the amino acid sequence common between the spindle-associated motif and the SGO motif (sequence shown in Suppl. Fig. S1A). (B) Fluorescence images of live cells of SGY8274 (*SGO1/SGO1-2GFP::TUB1/TUB1-RFP*) display dynamic localization patterns of Sgo1 at various stages of the cell cycle exhibiting co-localization with the spindle (Tub1) from metaphase till anaphase. The Sgo1 signals disappear in telophase. An untagged Sgo1 expressing strain SGY8276 (*TUB1/TUB1-RFP*) was used as a negative control. (C) *Left*, Sgo1-2GFP localization patterns in the wild type strain SGY8200 (*SGO1/SGO1-2GFP::TUB4/TUB4-MCHERRY*) in presence (−NOC) or absence (+NOC) of microtubules. *Right*, the spindle-like localization of Sgo1-2GFP in DMSO control (−NOC) *mad2Δ/Δ* cells in the strain SGY8248 (*mad2Δ/Δ::SGO1/SGO1-2GFP::TUB4/TUB4-MCHERRY*) disappears upon removal of the microtubules (+NOC). Scale bar ~5 μm.

### Sgo1 is not essential for viability in *C. albicans*

To test essentiality of Sgo1 in *C. albicans* during vegetative growth, we constructed a conditional mutant of *SGO1* using <u>g</u>ene <u>r</u>eplacement <u>a</u>nd <u>c</u>onditional <u>e</u>xpression (GRACE) strategy, wherein, one copy of *SGO1* was deleted and the remaining copy was placed under a regulatable promoter (*P*_*MET3*_). *MET3* promoter is repressed in the presence of <u>c</u>ysteine and <u>m</u>ethionine (CM) and is de-repressed in their absence. Whereas the cells with depletion of an essential protein CaSth1 (Prasad et al 2019) did not grow, we observed that Sgo1-depleted cells (*sgo1/P*_*MET3*_*SGO1*) did not lose viability (Suppl. Fig. S2A). This suggests that Sgo1, like in *S. cerevisiae*, is non-essential for mitotic growth in *C. albicans*. We then constructed a strain deleted for both the copies of *SGO1* (*sgo1Δ/Δ*) and verified the deletions by Southern hybridization (Suppl. Fig. S2B). We did not observe any phenotype of *sgo1Δ/Δ* cells under normal growth condition where the cells grew like wild type without any defect in chromosome missegregation or cell cycle arrest (Suppl. Fig. S2C). We observed a lower survival rate of *sgo1Δ/Δ* cells, when treated with anti-microtubule drug nocodazole (Suppl. Fig. S2D), a phenotype similar to Sc*sgo1-100* mutant (Indjeian et al 2005). The nocodazole sensitivity phenotype of *C. albicans sgo1* deletion mutant has been recently reported (Brimacombe et al 2019), while this manuscript was under preparation.

### Sgo1 shows dynamic localization patterns throughout the cell cycle, including on the mitotic spindle

To assess the functions of Sgo1 in the cell cycle of *C. albicans* we looked at the sub-cellular localization of Sgo1. For this, we fused one copy of *SGO1* with two tandem copies of Candida-optimized GFP (Gerami-Nejad et al 2001) at the carboxy (C)-terminus in the wild type strain SN148 to generate *SGO1/SGO1-2GFP* and on further marking of α tubulin with RFP to generate *SGO1/SGO1-2GFP::TUB1/TUB1-RFP* strains. We also constructed *SGO1/SGO1-2GFP::TUB4/TUB4-MCHERRY* strain expressing Sgo1-2GFP along with Tub4-mCherry that marks the spindle poles. We verified that Sgo1-2GFP was functional as *sgo1Δ/SGO1-2GFP::TUB4/TUB4-MCHERRY* strain with Sgo1-2GFP as sole source of Sgo1, did not show any sensitivity to nocodazole as compared to increased sensitivity of *sgo1Δ/Δ* mutant strain (*sgo1Δ/sgo1Δ::TUB4/TUB4-MCHERRY*) (Suppl. Fig. S2E).

Live cell analysis revealed that Sgo1-2GFP demonstrated dynamic localization patterns throughout the cell cycle except in telophase (Fig. 1B). In the unbudded G1/S cells, Sgo1-2GFP appeared as a dot-like signal close to MTOC indicative of kinetochore localization, as the *C. albicans* kinetochores earlier showed to remain clustered near MTOC (Thakur and Sanyal 2011, 2012). In *C. albicans* metaphase cells, Sgo1-2GFP could be seen as one dot or two dots near duplicated SPBs. Interestingly, with the advent of anaphase, Sgo1 shows a spindle-like localization, perhaps due to localization at the mitotic spindle, and persists there till late anaphase (telophase) when the signal disappears, presumably due to the spindle dis-assembly. The disappearance of Sgo1 signal at telophase could possibly be due to its dissociation and/or degradation (Fig. 1B). This persistent, dynamic and spindle localization pattern of Sgo1 is in contrast with what has been observed in other yeast species including *S. cerevisiae* and *S. pombe*.

Disappearance of Sgo1 with the spindle disassembly would suggest that its spindle localization might require intact MTs. To investigate that, we followed the Sgo1-2GFP staining with (−NOC) or without (+NOC) the MTs in the cells deleted for a SAC component *MAD2* to avoid SAC-mediated cell cycle arrest upon disruption of MTs (Fig. 1C). Although SPB separation was poor in the absence of MTs in *mad2Δ/Δ* mutant cells, we failed to observe a spindle-like localization of Sgo1 but observed either no or weak and diffused localization in nocodazole treated anaphase cells (Fig. 1C, +NOC) as compared to the cells in the presence of MTs (Fig. 1C, −NOC) suggesting that Sgo1 is indeed localized along the spindle MTs. To examine whether this localization is mediated by an association between the MTs and its MT binding motif that we identified through in-silico analysis (Fig. 1A), we expressed a mutant version of Sgo1 expressing Sgo1-MTΔ lacking the MT binding motif (from 258 to 358 residues) fused with GFP at the C-terminus (Suppl. Fig. S2F) as the sole source of Sgo1. However, we failed to detect any lack of spindle association of Sgo1 MTΔ-GFP as compared to Sgo1-GFP (Suppl. Fig. S2G). It is possible that Sgo1 uses some other domain or its interaction with some other MT binding protein(s) is required for its association with the MTs.

### Sgo1 has a role in timely spindle disassembly

Since we observed that Sgo1 associates with the mitotic spindle (Fig. 1B), we wished to investigate its significance. We failed to observe any gross abnormalities in spindle (marked by Tub1-GFP) morphology in the wild type and *sgo1Δ/Δ* strains (Suppl. Fig. S3A) negating any function of Sgo1 on the spindle assembly. However, when we counted those cells with SPB-SPB (Tub4-Tub4) distance > 6 μm, we observed 47% more cells in the *sgo1Δ/Δ* mutant than the wild type showed intact spindle suggesting a possible defect in the spindle disassembly in the mutant (Fig. 2A). Additionally, in support of the defect in spindle disassembly, we could observe that *sgo1Δ/Δ* mutant exhibited longer anaphase spindles as compared to the wild type cells. We used *mad2Δ/Δ* mutant, that showed wild type spindle lengths, to conclude that this spindle defect of *sgo1Δ/Δ* was distinct from its SAC related defect (see later results) at the centromeres (Fig. 2B). In HeLa cells, depletion of Sgo1 causes an increase in metaphase spindle length due to absence of tension between the sister chromatids (Salic et al 2004). To assess if the longer anaphase spindles in *C. albicans sgo1Δ/Δ* cells is a consequence of longer metaphase spindles, we analyzed the metaphase spindle lengths in the wild type and *sgo1Δ/Δ* cells but observed no significant difference between them (Suppl. Fig. S3B).

**Figure 2.**
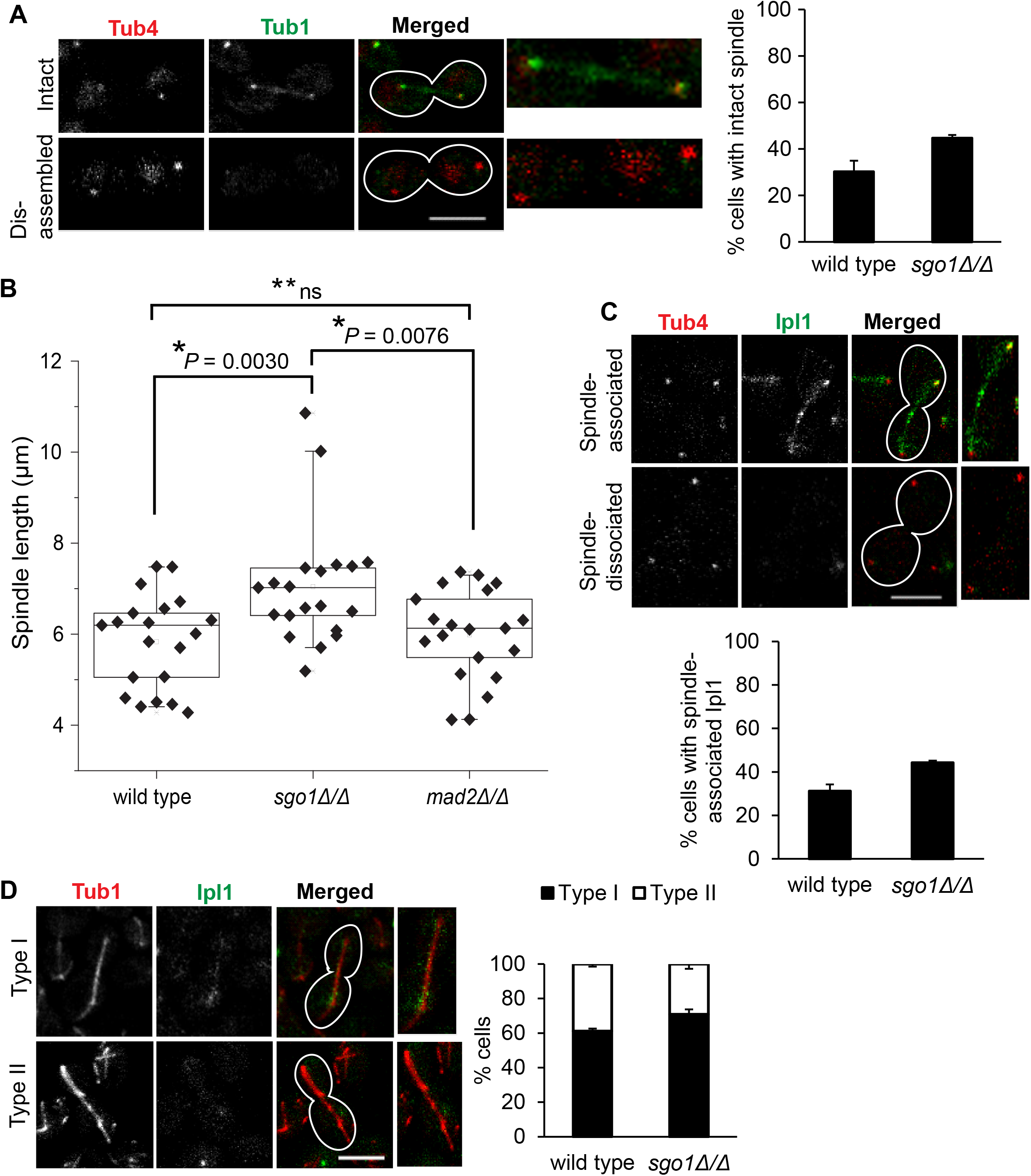
Sgo1 promotes timely spindle disassembly. (A) Live cell imaging of the wild type SGY8171 (*TUB1/TUB1-GFP::TUB4/TUB4-MCHERRY*) and *sgo1Δ/Δ* SGY8172 (*sgo1Δ/Δ::TUB1/TUB1-GFP::TUB4/TUB4-MCHERRY*) strains to visualize long anaphase spindle. Cells with SPB-SPB distance >6 μm were scored for presence (intact) or absence (disassembled) of such spindle. Histogram plot shows percentage of cells with intact spindle; the number of cells analysed, denoted here and in subsequent experiments as *n*, were at least 102 (*n* ≥ 102). (B) Analysis of spindle lengths in anaphase cells using the strains used in (A). Strain SGY8244 (*mad2Δ/Δ::TUB1/TUB1-GFP::TUB4/TUB4-MCHERRY)* was used as control; *n* = 20. (C) Live cell imaging of the wild type SGY8284 (*IPL1/IPL1-2GFP::TUB4/TUB4-MCHERRY*) and *sgo1Δ/Δ* SGY8290 (*sgo1Δ/Δ::IPL1/IPL1-2GFP::TUB4/TUB4-MCHERRY*) strains to assess presence (intact) or absence (dissociated) of Ipl1 in the cells with SPB-SPB distance > 6 μm. Histogram plot shows percentage of cells with intact spindle-like Ipl1 staining; *n* ≥ 105. (D) Live cell imaging of the wild type SGY8306 (*IPL1/IPL1-2GFP::TUB1/TUB1-RFP)* and *sgo1Δ/Δ* SGY8308 (*sgo1Δ/Δ::IPL1/IPL1-2GFP::TUB1/TUB1-RFP)* strains for simultaneous visualization of spindle and Ipl1. Type I and II show spindle (Tub1) with or without co-localization of Ipl1, respectively. Histogram plot depicts the distribution of type I and II among the cell population; *n* ≥ 103. Error bars here and in subsequent experiments denote standard error calculated from standard deviation obtained from three experimental replicates; asterisk (*) and (**ns) denote statistically significant (*P* < 0.05) and non-significant (*P* > 0.05) values, respectively calculated using two-tailed Student’s *t*-test. Scale bar ~5 μm.

Next, we sought to address how loss of Sgo1 might affect spindle disassembly. Since in *S. cerevisiae*, Ipl1 is required for spindle disassembly during late anaphase (Woodruff et al 2009), we argued that lack of Sgo1 in *C. albicans* may hinder normal timing of spindle disassembly by misregulating Ipl1 localization and/or its activity at the spindle. But we failed to observe any significant difference in gross intensity of Ipl1-2GFP along the spindle in anaphase cells (segregated SPBs, intact spindle) between wild type and *sgo1Δ/Δ* mutant (Suppl. Fig. S3C). However, similar to the observed intact long anaphase spindle (Fig. 2A), in the mutant 42% more cells over the wild type showed Ipl1 staining persistent along the spindle when SPB-SPB distance was more than 6 μm (Fig. 2C). In order to correlate the temporal dissociation of Ipl1 with spindle disassembly, we simultaneously monitored spindle (Tub1-RFP) and Ipl1. We scored cells harboring more than 6 μm spindle and assessed the presence or absence of Ipl1 on the spindle. The percentage of cells with Ipl1 staining on the spindle in the *sgo1Δ/Δ* mutant was comparable with the wild type (Fig. 2D, type I). We could not observe any early dissociation of Ipl1 from the spindle in the mutant cells. The percentage of cells with intact spindles but no Ipl1 on them was similar in the mutant and the wild type (Fig. 2D, type II). This suggests that in the mutant although Ipl1 resides on the longer spindle like wild type, it cannot promote spindle disassembly due to absence of Sgo1. These results indicate that the presence of Sgo1 at the spindle might be required for activity of Ipl1 at the spindle for its timely disassembly. We conclude that in *C. albicans*, Sgo1 displays a spindle disassembly function reminiscent of *S. cerevisiae* Ipl1, albeit does not perform it at least by mis-localizing Ipl1.

### Sgo1 associates with centromeric chromatin in cycling cells and in a SAC dependent way at the tensionless centromeres

ScSgo1 accumulates at the pericentromeres when the sister chromatids are not under tension whereas under unperturbed condition (presence of tension), it dissociates from the pericentromeres (Kiburz et al 2005, Nerusheva et al 2014). Since we observed nuclear localization of Sgo1 throughout the cell cycle in *C. albicans*, we wished to investigate whether Sgo1 associates with the centromeres even during unperturbed mitosis where the sister kinetochores are under tension. For this, one copy of the *SGO1* ORF was fused with Tandem Affinity Purification (TAP) tag at the C-terminus in the wild type SN148 strain to construct the *SGO1/SGO1-TAP*. Western blot analyses and nocodazole sensitivity assay confirmed the expression of Sgo1-TAP and its functionality, respectively (Suppl. Fig. S4 and S2D). Sgo1-TAP ChIP using anti-Protein A antibodies revealed that Sgo1 is indeed significantly associated with the centromeres in presence or absence of nocodazole (Fig. 3A) suggesting that Sgo1 might have conserved its tension-sensing function in *C. albicans*. The localization of Sgo1 at the centromeres even in the presence of tension (Fig. 3A) argued for additional role(s) of Sgo1 in *C. albicans*.

**Figure 3.**
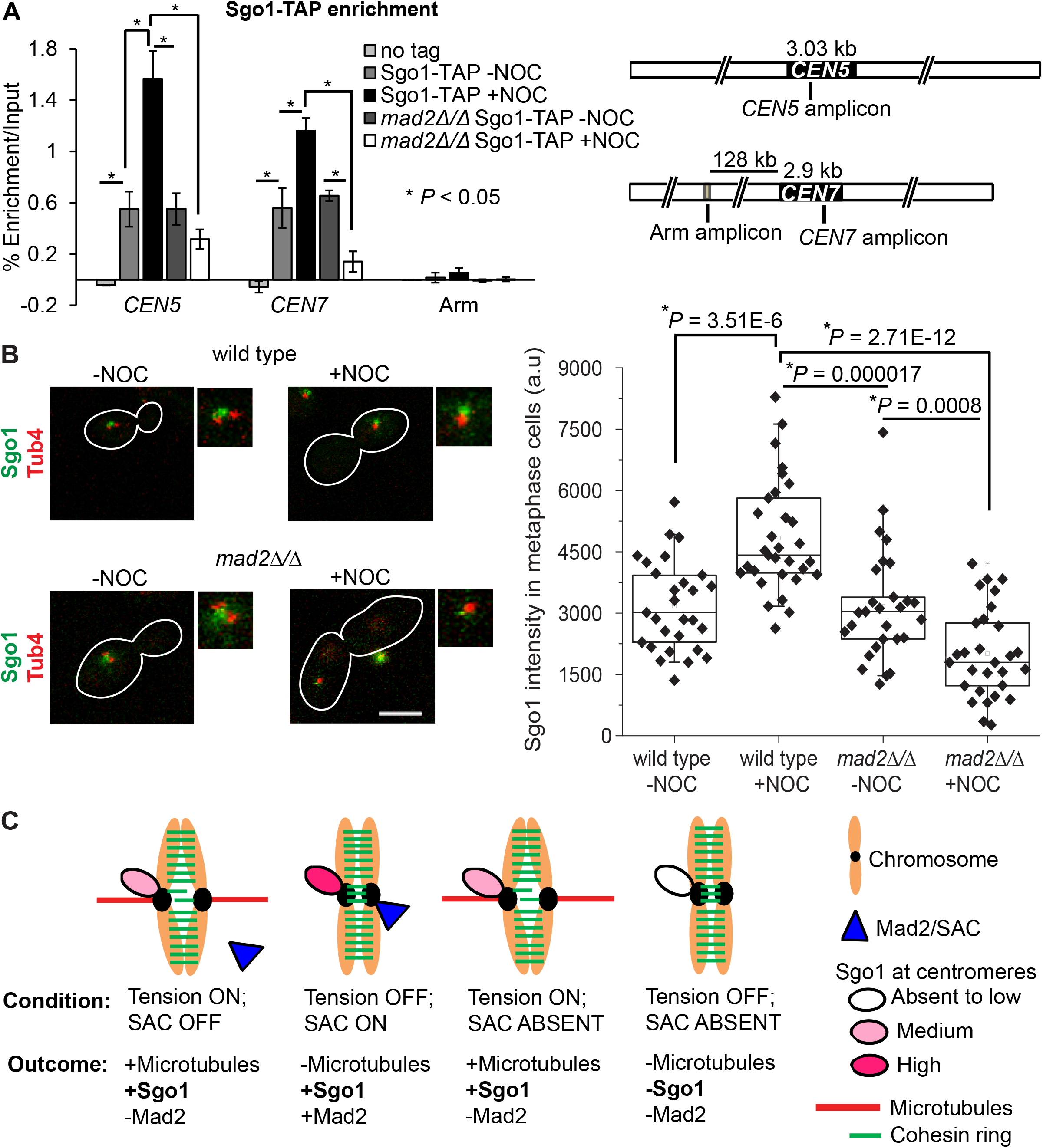
Mad2-driven increase in centromeric Sgo1 levels upon loss of tension. (A) Sgo1-ChIP assays were performed either in presence or absence of nocodazole in SGY8038 (*SGO1/SGO1-TAP*) and SGY8130 (*mad2Δ/Δ::SGO1/SGO1-TAP*) to assess the binding of Sgo1 at centromeres and at a non-centromeric locus (128 kb distal to *CEN7*). SN148 (wild type) was used as no tag control. Relative positions of the amplicons corresponding to centromeres and arm regions on chromosome 5 and 7 are shown below. (B) Box plot depicting background corrected fluorescence intensities of Sgo1-2GFP signals in the wild type SGY8156 (*SGO1/SGO1-2GFP*) and *mad2Δ/Δ* SGY8248 (*mad2Δ/Δ::SGO1/SGO1-2GFP::TUB4/TUB4-MCHERRY*) strains in absence or presence of nocodazole; *n* ≥ 28. (C) Schematic of Sgo1 and Mad2 association with the centromeres under various conditions.

Since an absence of tension causes SAC localization at the centromeres, we sought to examine whether in the absence of tension, Sgo1 is enriched at the centromeres in a SAC-dependent manner. Sgo1-TAP ChIP in *mad2Δ/Δ* mutant strain (*mad2Δ/Δ::SGO1/SGO1-TAP*), in the presence or absence of microtubules, was performed. A significant drop in Sgo1 localization at the centromeres in the absence of Mad2 as compared to its presence (Fig. 3A) confirmed that Sgo1 enrichment at the centromeres is Mad2-dependent at metaphase. However, Sgo1 localization, albeit at lesser extent, at the centromeres in the cycling cells is Mad2 independent (compare wild type and *mad2Δ/Δ*, −NOC, Fig. 3A). Further, Sgo1-2GFP intensity in the wild type and *mad2Δ/Δ* metaphase stage cells in the presence (−NOC) or absence (+NOC) of microtubules (Fig. 3B) was found to be consistent with the ChIP data suggesting that the alterations in Sgo1 level is not due to fixation of the cells with formaldehyde. Based on these observations, we conclude that Sgo1 is associated with centromeres under normal condition in a Mad-independent manner but its recruitment at the centromeres under tensionless condition requires Mad2 (Fig. 3C).

### Absence of Sgo1 relieves SAC mediated G2/M arrest caused by kinetochore-MT attachment defect

The enrichment of Sgo1 at the microtubule unattached kinetochore in a SAC-dependent way prompted us to investigate if this protein has any role in SAC function. If Sgo1 is required for SAC function, then absence of this protein should relieve SAC-mediated G2/M arrest caused due to disruption of KT-MT attachment by inactivation of kinetochore. To examine this, we used a CM repressible outer kinetochore mutant *dam1* that was shown to arrest at G2/M stage in a SAC-dependent way resulting in large-bud cells with unsegregated nuclei (Thakur and Sanyal 2011). An arrest-relieved phenotype was expected in Dam1-depleted cells with no SAC or Sgo1 function. Analysis of large bud population and DAPI segregation showed that the wild type, *sgo1Δ/Δ* and *mad2Δ/Δ* mutant showed no significant arrest phenotype when Dam1 was expressed (-CM) (Fig. 4A, 1^st^, 2^nd^ and 3^rd^ rows, respectively). As expected Dam1 depleted cells showed high frequency (97%, 6 h, Fig. 4A, right panel) of G2/M arrest phenotype and removal of SAC (Mad2) in those cells caused a precipitous drop (23%, 6 h) in that phenotype indicating relief from the arrest. Interestingly, removal of Sgo1 in the Dam1 depleted cells also caused similar arrest-relieved phenotype albeit to a lesser extent (54%, 6 h). FACS analysis confirmed that Dam1 depletion in the absence of either Mad2 or Sgo1 leads to a higher accumulation of aneuploid cells as compared to when both Mad2 and Sgo1 are present presumably as a result of bypass of the arrest without proper chromosome segregation (Fig. 4B). Bypass of the cell cycle arrest in presence of defect in KT-MT attachment due to compromised SAC function can cause genomic instability and consequent drop in cell viability. Indeed, the cells devoid of both Dam1 and Mad2 or Sgo1 displayed a more dramatic loss in viability than the cells devoid of Dam1 alone (Fig. 4C). To support these observations, we recreated defective KT-MT attachment by removing microtubules by treating cells with anti-microtubule drug nocodazole for increasing duration and *sgo1Δ/Δ* cells were analyzed for large bud (mitotic) arrest and viability along with the control cells. We observed a sharp decrease in viability from 60 to 0% after 2 to 8 h of nocodazole treatment which was similar to the phenotype of the SAC mutant, *mad2Δ/Δ* (Fig. 4D and Suppl. Fig. S5A). To confirm the SAC function of Sgo1, we examined if *sgo1Δ/Δ* cells could bypass metaphase arrest imposed by nocodazole. Strikingly, while the wild type cells became elongated due to arrest, the *sgo1Δ/Δ* cells instead showed mostly multi-budded phenotype, similar to the *mad2Δ/Δ* cells, suggesting progress through the cell cycle (Suppl. Fig. S5B, 4, 6 and 8 h). These results altogether support that Sgo1 promotes SAC function in *C. albicans*.

**Figure 4.**
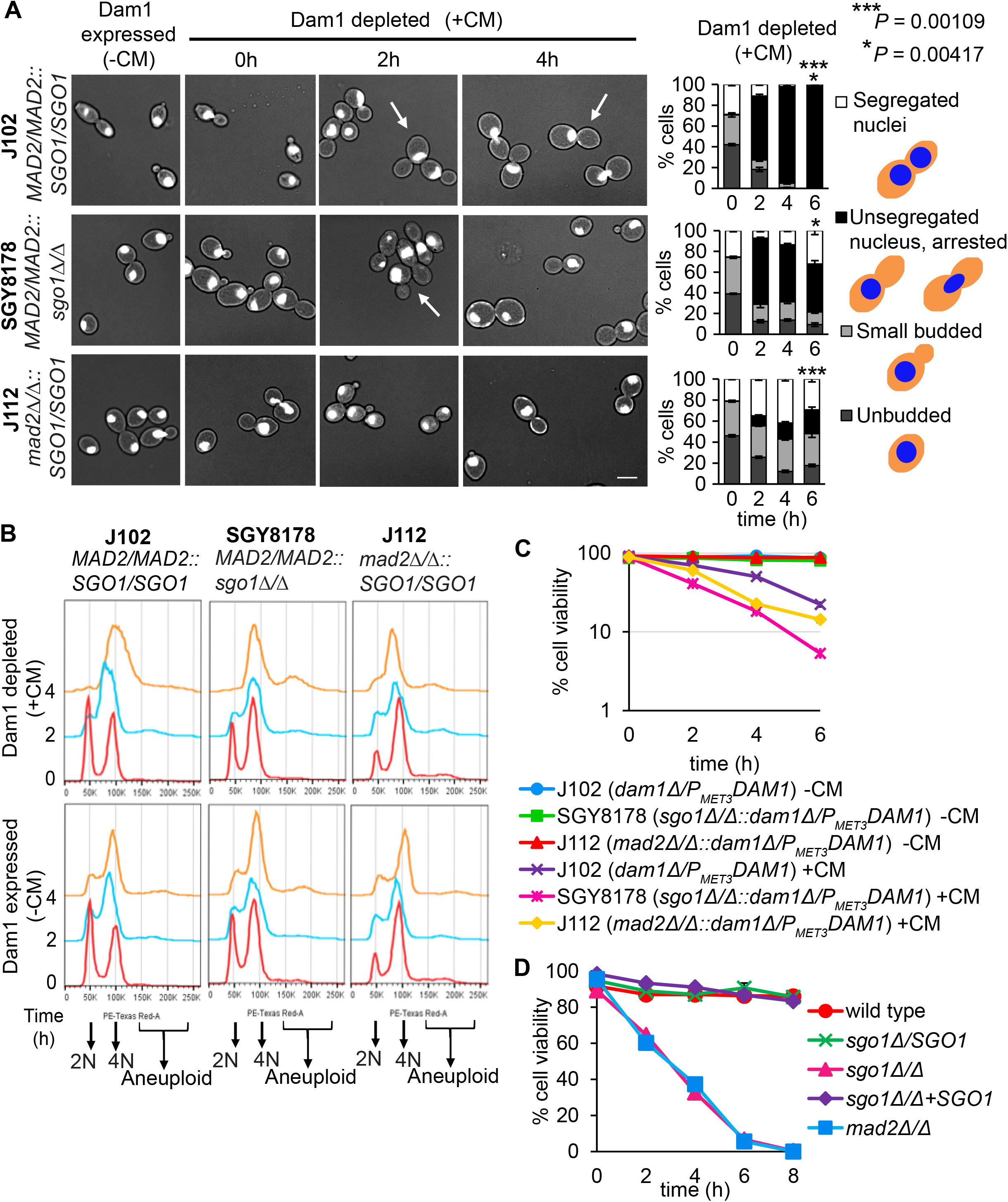
Absence of Sgo1 relieves Mad2-mediated G2/M arrest. (A) Strains J102 (*dam1Δ/P*_*MET3*_*DAM1*), SGY8178 (*sgo1Δ/Δ::dam1Δ/P*_*MET3*_*DAM1*) and J112 (*mad2Δ/Δ::dam1Δ/P*_*MET3*_*DAM1*) were analysed for budding index and nuclear segregation under permissive (-CM) or non-permissive condition (+CM) for Dam1 expression at the indicated time intervals. *Left*, representative images. *Right*, distribution of the representative phenotypes when Dam1 is depleted for indicated durations; *n* ≥ 98, ‘unsegregated nucleus, arrested’ population at 6 h was compared between the strains to calculate *P* values. * denotes comparison between J102 and SGY8178; *** denotes comparison between J102 and J112. Scale bar ~5 μm. (B) The same strains were analysed by FACS following indicated time of growth under Dam1 depleted or expressed conditions. (C) Viability assay of the above strains under Dam1 expressed (-CM) or depleted (+CM) conditions. (D) Viability assay of wild type SGY8110 (*SGO1/SGO1*), SGY8112 (*sgo1Δ/SGO1)*, SGY8114 (*sgo1Δ/Δ*), SGY8116 (*sgo1Δ/Δ:SGO1)* and J110 (*mad2Δ/Δ*) strains, treated with 20 μg/ml nocodazole for the indicated time, harvested, washed and plated on SD agar plates without nocodazole. The colonies appeared on the plates were counted to estimate the viability.

To confirm that the *sgo1Δ/Δ* cells are sensitive to perturbation in kinetochore-microtubule related process but are not deficient in DNA damage response (DDR), we assessed the sensitivity of *sgo1Δ/Δ* cells to DNA damaging drug hydroxyurea (HU). DDR protein CaMec1 was used as a control (Legrand et al 2011). We observed that compared to *mec1Δ/Δ*, *sgo1Δ/Δ* cells did not show any sensitivity to 20 mM HU (Suppl. Fig. S5C), suggesting that Sgo1 functions specifically during G2/M arrest mediated by SAC and not during S-phase arrest mediated by the DDR checkpoint.

### Sgo1 maintains SAC proteins at the kinetochore during prolonged SAC-mediated arrest

Since, in *C. albicans*, Sgo1 exhibits SAC function (Fig. 4) where Sgo1 and Mad2 may function in tandem to ensure efficient checkpoint arrest, we reasoned that these two proteins may interact directly or indirectly. To investigate if localization of Mad2 at the unattached kinetochore depends on Sgo1, we tagged Mad2 with 3HA at the C-terminus in wild type and *sgo1Δ/Δ* strains, harboring Cse4-GFP as the centromere marker. Absence of Sgo1 did not affect the expression of Mad2 (Suppl. Fig. S6A). We then performed the chromatin spread assay after treating these strains with nocodazole for increasing duration. Following 2 h of treatment, there was no difference in Mad2 co-localization with Cse4-GFP between wild type and *sgo1Δ/Δ* strains (Fig. 5A). However, when we prolonged the duration of nocodazole-mediated arrest to 4 h, strikingly *sgo1Δ/Δ* cells displayed either loss of tight knit centromeric Mad2 signal or appearance of some weak diffuse signal (Fig. 5A) in 48% more number of spreads compared to the wild type. This suggests that although Sgo1 is not essential for recruitment of Mad2 at the kinetochore, it becomes crucial for maintenance of Mad2 during prolonged SAC mediated cell cycle arrest which is consistent with abrogation of this arrest in *sgo1Δ/Δ* (Fig.4 and Suppl. Fig. S5A, B). To investigate if Sgo1 function is specific to Mad2 or to other SAC proteins as well, we assessed the localization of another SAC protein, Bub1 which is known to act upstream of Mad2 in budding yeast (Chen et al 1999; London and Biggins, 2014; Moyle et al 2014). Like Mad2, a tight knit Bub1 signal next to SPB indicated its centromeric localization (Fig. 5B, type I), whereas a diffuse or no signal indicated its dislodgement from the centromeres (Fig. 5B, type II and III). We observed, that the Bub1 localization at the centromeres in cycling wild type and *sgo1Δ/Δ* cells is comparable (Fig. 5B, −NOC, top panel). However, in response to the metaphase arrest following 2 h of nocodazole treatment, the localization of Bub1 at the centromeres appears reduced, albeit to a lesser extent, in *sgo1Δ/Δ* as compared to wild type (Fig. 5B, +NOC, bottom panel). Following 4 h of nocodazole treatment, the percentage of cells displaying Bub1 dissociated from the centromeres increases drastically, suggesting that Sgo1 functions in the maintenance of more than one SAC proteins at the centromeres during G2/M arrest.

**Figure 5.**
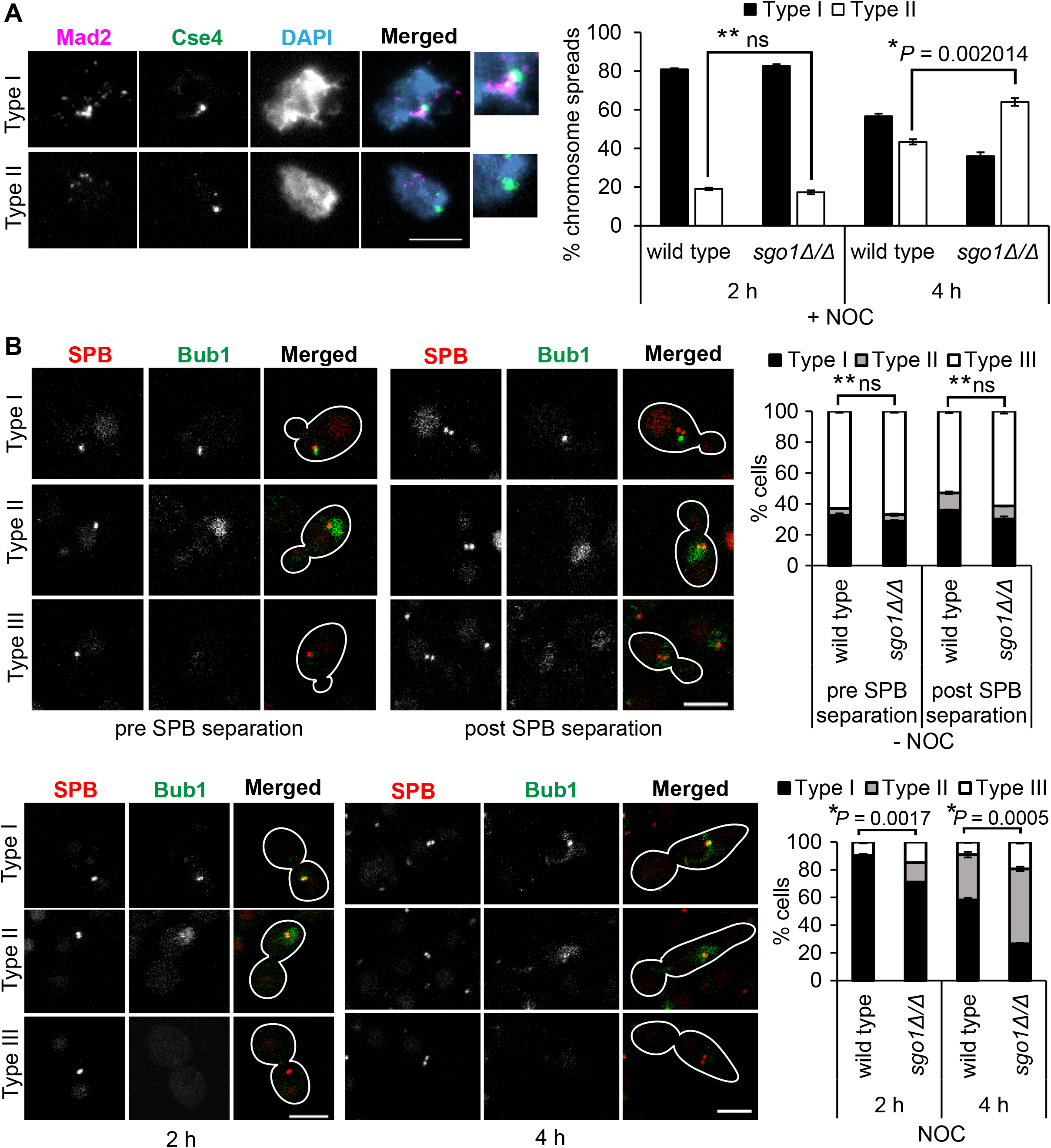
Sgo1 maintains spindle assembly surveillance components Mad2 and Bub1 at centromeres during prolonged checkpoint arrest. (A) Representative images of chromatin spreads showing the localization of Mad2-3HA in presence of 20 μg/ml nocodazole for indicated time durations using the wild type SGY8301 (*CSE4/CSE4-GFP-CSE4::MAD2/MAD2-3HA*) and *sgo1Δ/Δ* SGY8302 (*sgo1Δ/Δ*::*CSE4/CSE4-GFP-CSE4::MAD2/MAD2-3HA*) strains. Type I denotes localization of Mad2 at the centromeres whereas type II indicates no or weak and diffuse localization of Mad2; *n* ≥ 50. (B) Representative images of cells showing Bub1-GFP localization using the wild type SGY8326 (*SGO1/SGO1::BUB1/BUB1-GFP::TUB4/TUB4-MCHERRY*) and *sgo1Δ/Δ* SGY8328 (*sgo1Δ/Δ::BUB1/BUB1-GFP::TUB4/TUB4-MCHERRY)* strains treated without (top panel) or with (bottom panel) 20 μg/ml nocodazole for indicated stages and time points, respectively. The distribution of type I-III categories are shown on right of each panel; *n* ≥ 96, type I population was compared between the strains to calculate *P* values. Scale bar ~ 5 μm.

A report in fission yeast has revealed a Mad2-independent role of Sgo1 in delaying anaphase onset (Meadows et al 2017). If Sgo1 in *C. albicans* possesses any Mad2-independent function at the centromeres, then *sgo1 mad2* double mutant is expected to show synergistic effect upon nocodazole treatment. To assess this, we constructed *sgo1 mad2* double mutant and subjected it to nocodazole mediated metaphase arrest. Large budded or elongated cells with undivided DAPI was considered as arrested cells whereas such cells with segregated DAPI and multibudded cells were considered to have bypassed the arrest. We observed that the extent of bypass of arrest in *sgo1 mad2* double mutant was slightly more as compared to either *sgo1* or *mad2* single mutant (Suppl. Fig. S6B, bottom panel, compare the pair of bars for the mutants with *P* value <0.01). Consistent to more bypass of arrest, a higher drop in cell viability was observed in *sgo1 mad2* double mutant (Suppl. Fig. S6C) than in *sgo1* or *mad2* single mutant suggesting that Sgo1 might have additional function(s) at the centromeres in response to metaphase arrest, which perhaps is independent of Mad2.

### Sgo1 ensures chromosome biorientation and disjunction

ScSgo1 is required for de novo amphitelic attachment of the sister chromatids with the microtubules emanating from opposite spindle poles and therefore, the protein becomes essential during recovery from anti-microtubule drug treatment (Indjeian and Murray 2007). To examine if Sgo1 has a similar role in sister chromatids bi-orientation in *C. albicans*, the wild type and the mutant (*sgo1Δ/Δ*) strains harboring an array of *tet* operators integrated immediately adjacent to *CEN7* on one copy of chromosome 7 and expressing TetR-GFP fusion protein (Burrack et al, 2013) were used. The spindle poles (SPBs) were marked with Tub4-mcherry in these strains. For biorientation assay, we treated both the wild type and the mutant strains with nocodazole to disassemble the spindle, and then released them into nocodazole-free media to allow for de novo reassembly of the mitotic spindle (Suppl. Fig. S7A). Biorientation was scored as two GFP dots each proximal to two separated SPBs resulting from amphitelic attachment (Fig. 6, type I, left panel). In case of erroneous syntelic and monotelic (monooriented) attachments, two or one GFP dot(s) (depending upon the presence of cohesion between the sister chromatids) remained proximal to one of the two separated SPBs (Fig. 6, type II, left panel). We observed that following spindle recovery, in the wild type almost 83% of the cells achieved sister chromatid biorientation whereas only 47% could achieve so in *sgo1Δ/Δ* cells (Fig. 6, right panel) suggesting that the function of Sgo1 in chromosome biorientation is conserved in *C. albicans*.

**Figure 6.**
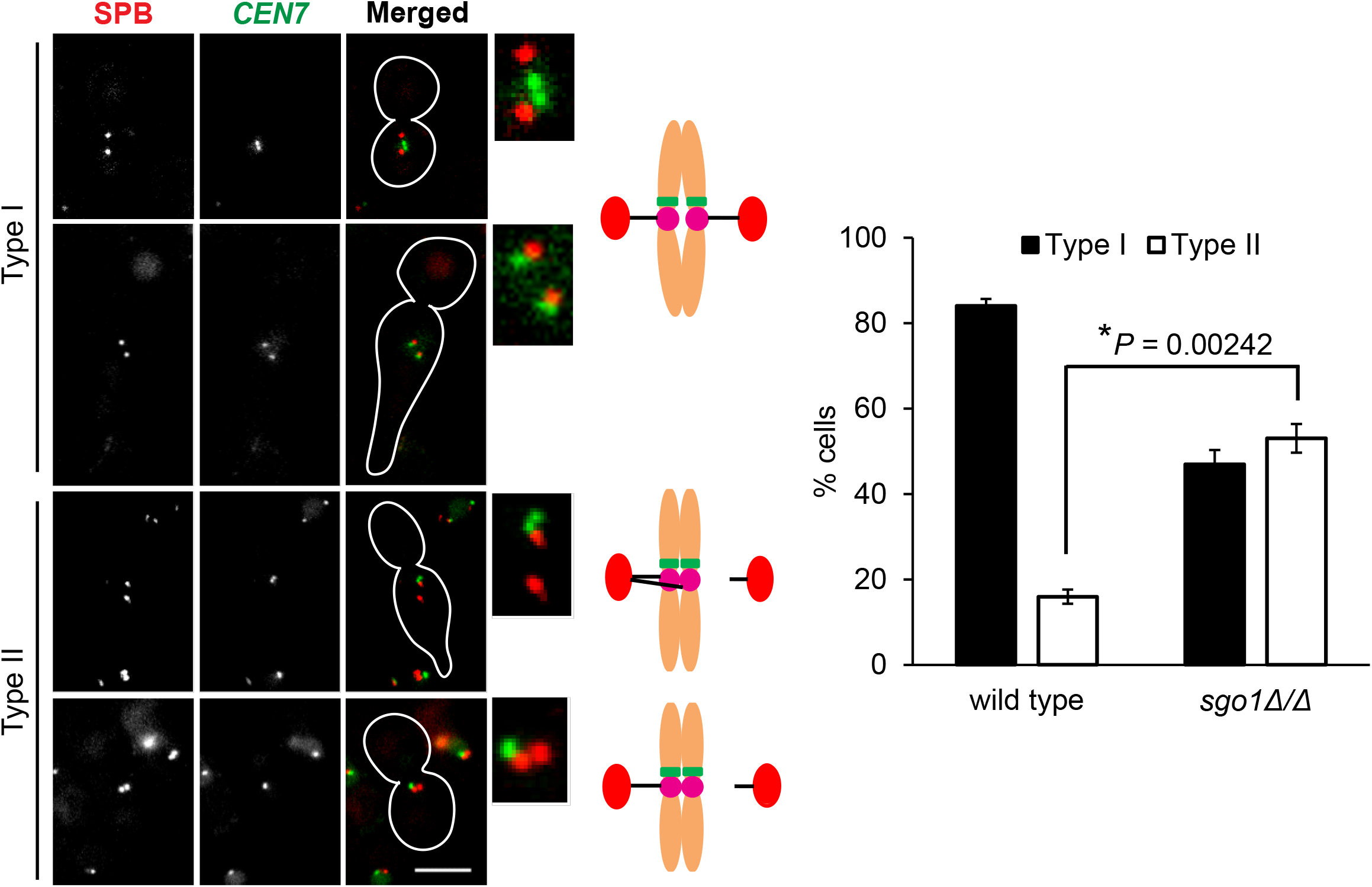
Sgo1 ensures chromosome biorientation. The cells from the wild type SGY8154 (*TetO-CEN7/TetR-GFP*::*TUB4/TUB4-MCHERRY*) and *sgo1Δ/Δ* SGY8165 (*sgo1Δ/Δ::TetO-CEN7/TetR-GFP*::*TUB4/TUB4-MCHERRY*) strains were treated with 20 μg/ml nocodazole for 90 min and were released into drug-free media for 15 min before they were imaged. Representative images of the cells showing bioriented (type I) or monooriented (type II) sister chromatids. In both these strains, SPBs were marked with Tub4-mcherry and one copy of chromosome 7 was marked with TetO-GFP-TetR (see text). Scale bar ~5 μm. Histogram plot depicts distribution of type I and II categories among the cell population of the indicated strains; *n* ≥ 186.

Errors in chromosome biorientation during spindle recovery can lead to chromosome non-disjunction, generation of aneuploidy and loss of cell viability. Since we observed *sgo1Δ/Δ* cells lose viability over the wild type while recovering from spindle poison (Fig 4D), we examined the chromosome segregation in these cells following full spindle recovery. We observed that in wild type strain around 86% of the cells showed proper chromosome disjunction, whereas in *sgo1Δ/Δ* strain the same dropped to 43% (Suppl. Fig S7B). This further confirmed that during establishment of de novo KT-MT attachment Sgo1 is required to promote sister chromatin biorientation to ensure accurate chromosome segregation.

### Sgo1 is required for condensin recruitment and a sustained Ipl1 localization at centromeres

Since we could detect Sgo1 at the centromeres in the unperturbed cycling cells (Fig. 3A), we wished to investigate if, besides SAC related function, there is any other role of this protein at the centromeres in *C. albicans*. In higher eukaryotes, Sgo1 is involved in the maintenance of pericentromeric cohesin during mitotic prophase pathway which removes the arm cohesion (Hauf et al 2005, Kueng et al 2006, Tang et al 2006, Shintomi and Hirano 2009). Additionally, across sexually reproducing organisms, Sgo1 protects centromeric cohesin during meiosis I (reviewed in Marston 2015). Therefore, we wished to test whether Sgo1 has any role in sister chromatid cohesion during mitosis in *C. albicans*. We performed the <u>s</u>ister <u>c</u>hromatid <u>c</u>ohesion (SCC) assay using the strains used for the biorientation assay. Both the wild type and *sgo1Δ/Δ* strains were treated with 20 μg/ml nocodazole to remove the microtubule-based pulling force. Under this tension-less condition, in the cells with undivided DAPI mass, the cohered sister-*CEN7-*GFP dots will appear as a single dot (Fig. 7A, +NOC, type I) due to diffraction limit, whereas non-cohered sisters will appear as two separate dots (Fig. 7A, +NOC, type II). We failed to observe any significant percentage of cells with separated GFP dots both in the wild type and in the *sgo1Δ/Δ* cells suggesting loss of Sgo1 does not compromise sister chromatin cohesion (Fig. 7A, +NOC, right panel). This result is consistent with our observations that *sgo1Δ/Δ* cells do not exhibit chromosome missegregation and loss in cell viability under unperturbed conditions (Suppl. Fig. S2C). Interestingly, in the cycling cells at the metaphase stage (assessed by large-budded cells with undivided DAPI mass), where the sister kinetochores are under microtubule-based pulling force, we observed a greater percentage (36%) of the mutant cells displaying separated GFP dots as compared to the wild type (12%) (Fig. 7A, −NOC, right panel). This indicates that the cohesive force between the sisters is less in the *sgo1* mutant although the SCC assay could not detect any cohesin defect in the mutant. To reconcile these observations, it can be argued, that the absence of Sgo1 reduces the tensile strength of chromatin as reported earlier in *S. cerevisiae* (Peplowska et al 2014) and hence the chromatin becomes more stretchable causing the GFP dots to split asunder more frequently upon microtubule mediated pulling force, given that the lack of Sgo1 does not perturb the bi-orientation in cycling cells.

**Figure 7.**
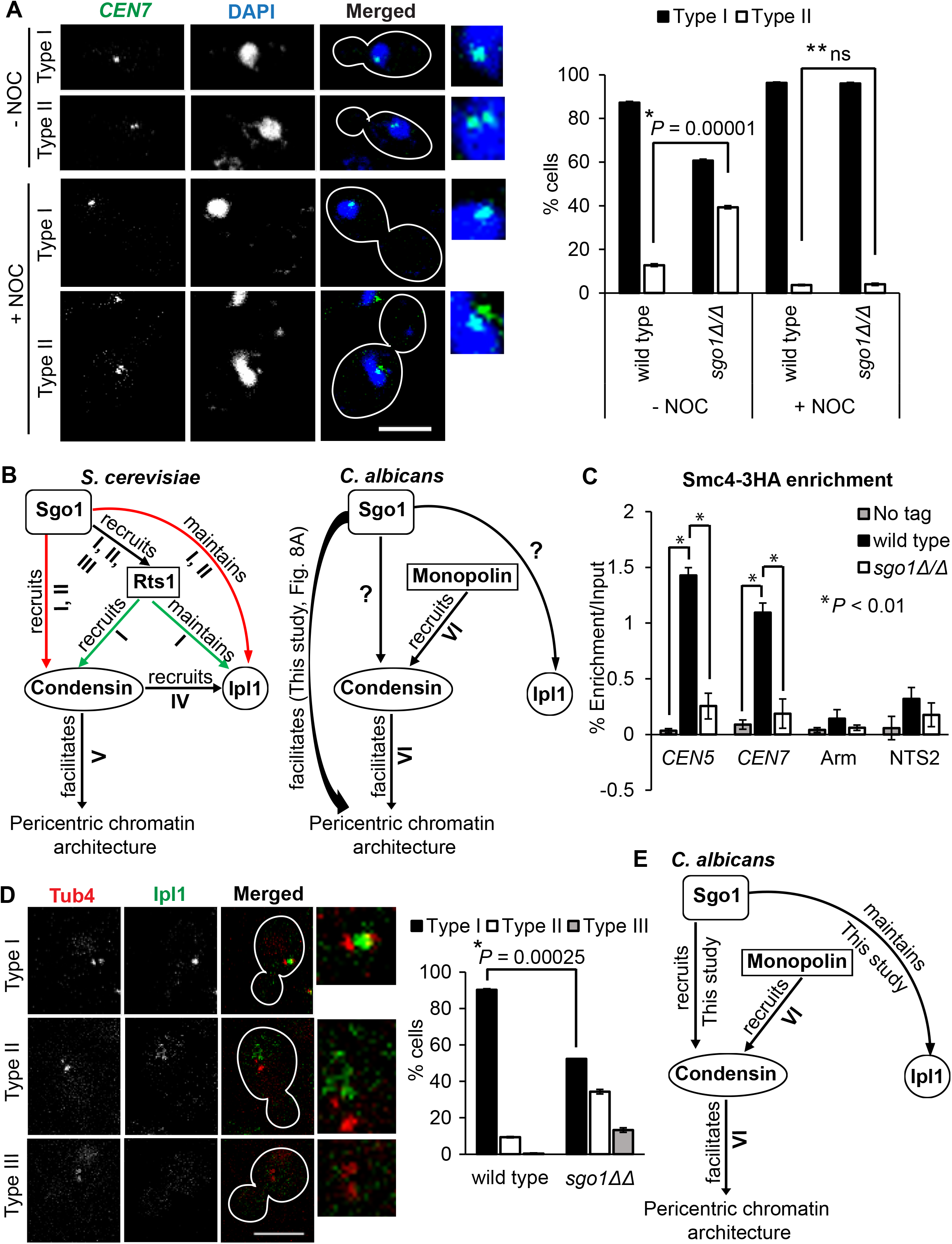
Loss of Sgo1 perturbs pericentromeric chromatin by affecting localization of condensin and maintenance of aurora B kinase Ipl1 at centromeres. (A) Live cell imaging of the strains YJB13024 (wild type) and SGY8121 (*sgo1Δ/Δ*) harboring heterozygous *CEN7*-GFP in presence or absence of 20 μg/ml nocodazole. Type I and II denote unseparated and separated sister centromeres visualized as 1 and 2 GFP dot(s), respectively; *n* ≥ 87. (B) Model of putative functions of Sgo1 in *C. albicans* based on its functions with respect to condensin and aurora B kinase (Ipl1) in *S. cerevisiae*. Green and red arrows depict Rts1 dependent and independent recruitment, respectively. I, Verzijlbergen et al 2014; II, Peplowska et al 2014; III, Eshleman et al 2014; IV, Li et al 2011; V, Stephens et al 2011; VI, Burrack et al 2013. (C) ChIP assays with anti-HA antibodies to detect Smc4-3HA binding at the centromeres, rDNA locus (NTS2) and arm region using the wild type SGY8238 (*SMC4/SMC4-3HA*) and *sgo1Δ/Δ* SGY8227 (*sgo1Δ/Δ::SMC4/SMC4-3HA*) strains. No tag strain SN148 (*SMC4/SMC4*) was used as a control. (D) Centromeric localization of Ipl1 in pre-anaphase cells (<2 μm spindle) using wild type SGY8284 (*IPL1/IPL1-2GFP::TUB4/TUB4-MCHERRY*) and *sgo1Δ/Δ* SGY8290 (*sgo1Δ/Δ::IPL1/IPL1-2GFP::TUB4/TUB4-MCHERRY*) strains. Type I, II and III depict centromeric, diffuse and no signals of Ipl1, respectively; *n* ≥ 101. (E) Working model depicting functions of Sgo1 in localization of condensin and maintenance of Ipl1 in *C. albicans* revealed in this study. Scale bar ~5 μm.

In *C. albicans*, abrogation of centromeric association of condensin leads to decreased chromosome stiffness as metaphase cells show an increased distance between sister chromatids as compared to the wild type (Burrack et al 2013). Similar to this result, since we observed an increased percentage of metaphase cells with separated sister chromatids in *sgo1Δ/Δ* as compared to the wild type (Fig. 7A), we argued, that Sgo1 perhaps conserved the function of condensin recruitment in *C. albicans* (Fig. 7B). To investigate this, we first tagged a condensin subunit Smc4 with 3HA at the C-terminus and verified that its expression is comparable in wild type and *sgo1Δ/Δ* cells (Fig. S8A). To assess the centromeric localization, we performed Smc4-ChIP and observed that in the absence of Sgo1, centromeric localization of Smc4 significantly dropped as compared to the wild-type (Fig. 7C). However, as a negative control, we failed to detect any significant difference in Smc4 recruitment at the Non-transcribed spacer (NTS2) region of rDNA locus. Together, above results suggest that in *C. albicans*, Sgo1 has retained the function of centromeric recruitment of condensin and maintenance of chromosome stiffness (Fig. 7E).

Since in *S. cerevisiae* condensin has been implicated in centromeric localization of aurora B kinase, Ipl1 (Li et al 2011) and Sgo1 has been shown to be essential for Ipl1 maintenance at the centromeres (Verzijlbergen et al 2014, Peplowska et al 2014), we expected *C. albicans* Sgo1 to be involved in regulating Ipl1 localization (Fig. 7B). To assess this, we followed Ipl1-2GFP localization in wild type and *sgo1Δ/Δ* with respect to SPBs (Tub4-mcherry) in live cells. The localization of Ipl1 as tight knit dot next to one SPB or in between two closely separated SPBs was considered its centromeric localization (Fig. 7D, type I, left panel), as the centromeres remain clustered next to SPB in *C. albicans* throughout the cell cycle (Sanyal and Carbon 2002; Roy et al 2011, Thakur and Sanyal 2011). Early at the cell cycle when SPBs are not separated, we did not observe any difference in localization of Ipl1 at the centromeres between wild type and *sgo1Δ/Δ* cells (Suppl. Fig. S8B). However, in the cells with separated SPBs (SPB-SPB distance <2 μm), the wild type maintained the centromeric signal of Ipl1 in majority of the cells but a considerable fraction of *sgo1Δ/Δ* cells displayed either diffuse (type II) or no signal (type III) of Ipl1 (Fig. 7D, right panel) indicating its dislodgement from the centromeres. These observations suggest that, like in *S. cerevisiae*, Sgo1 is not required for initial recruitment of Ipl1 at the centromeres, but becomes crucial for its maintenance at those loci (Fig. 7E).

## Discussion

In concordance with the name of “guardian spirit” in Japanese, shugoshin family of proteins play key roles in regulating several events of eukaryotic chromosome segregation and thus maintain the ploidy. Importantly, without harboring any enzymatic function, shugoshin regulates those events by acting as a molecular adapter by recruiting myriad proteins as effectors at various subcellular locales in a timely manner. Although certain functions of shugoshin are conserved across eukaryotes, a considerable variation of functions have also been observed in a species-specific manner. In this context, since *C. albicans* remarkably uses ‘aneuploidy in moderation’ as a tool to strive over the stress conditions, it was tempting to investigate how shugoshin, in general, functions in this organism with, if any, functional rewiring.

### Single shugoshin protein in *C. albicans*

Since shugoshin was first identified in flies as MEI-S332 (Kerrebrock et al 1992), this protein exists in nearly all eukaryotes as a single protein or as two paralogues. While in flies and budding yeast, a single shugoshin is present, in fission yeast and in vertebrates two paralogues are present. Through in-silico analysis, in *C. albicans* we found a single shugoshin protein (Suppl. Fig. 1). In organisms with two paralogues, each protein has separate and overlapping functions suggesting that the separation of functions occurred late during evolution. In fact, it is believed that the centromeric cohesion protection function of shugoshin has been acquired later, from its function in sister chromatid bi-orientation (Vanoosthuyse et al 2007, Kawashima et al 2007). In the absence of ‘prophase pathway’ and meiosis in *C. albicans*, the requirement of centromeric cohesin protection function can be dispensable, and therefore this organism might have evolved with a single shugoshin protein having bi-orientation function. However, in this context, the scenario in *S. cerevisiae*, due to its unique point centromere, is perhaps non-canonical as there is a single shugoshin which performs both the functions (Kitajima et al 2004, Indjeian et al 2005, Indjeian and Murray 2007, Peplowska et al 2014).

### Spindle localization of shugoshin and function in spindle disassembly

Till date, among the species, shugoshin has been found on the chromatin at the centromeres/pericentromeres (Kitajima et al 2004, Kiburz et al 2005) and in the vicinity of telomeres (Vanoosthuyse et al 2007, Tashiro et al 2016). Its association with the chromosomal arms cannot be ruled out given its function on arm condensation in budding yeast (Kruitwagen et al 2018) and in the prophase checkpoint in *C. elegans* (Bohr et al 2018). Our finding that shugoshin in *C. albicans* is localized along the microtubule spindle till late anaphase (Fig. 1) is intriguing. Earlier it has been shown that in *Xenopus* and in *Mus musculus*, Sgo1 can bind to the microtubules and promote microtubule polymerization under an in vitro condition (Salic et al 2004). In vivo implication of Sgo1-microtubule interaction has also been demonstrated in HeLa cells where the microtubules are destabilized in absence of Sgo1 (Salic et al 2004). Furthermore, in humans a spliced variant of Sgo1 (sSgo1) has been found to be localized at the mitotic spindle and centrosomes, a function that has been implicated in regulation of spindle pole duplication and cohesion between the mother and daughter centrioles (Wang et al 2006, Wang et al 2008). While these studies in vertebrates account for a microtubule stabilizing role of shugoshin, in contrast our results indicate that in *C. albicans*, microtubule localized Sgo1 is involved in timely spindle disassembly (Fig. 2). This difference might be subjected to unique prolonged spindle association of CaSgo1 from metaphase till late anaphase (Fig. 1) which is not observed in vertebrates (Salic et al 2004). It is important to address the plausible mechanism by which CaSgo1 might promote spindle disassembly. Since Ipl1/Aurora B kinase/Ark1 has been shown to promote spindle disassembly in budding yeast (Buvelot et al 2003) and fission yeast (Flor-Parra et al 2018) and shugoshin as adapter recruits aurora B (CPC) at the centromeres in various organisms including budding yeast (Peplowska et al 2014), fission yeast (Kawashima et al 2007, Vanoosthuyse et al 2007), *Xenopus* (Rivera and Losada 2009, Shintomi and Hirano 2009) and Hela cells (Tsukahara et al 2010), we argued that CaSgo1 might promote spindle disassembly by facilitating Ipl1 localization at the spindle. However, we failed to detect any mislocalization of Ipl1 in *sgo1Δ/Δ* cells, rather more cells with long spindles decorated with Ipl1 were observed (Fig. 2). This indicates perhaps Ipl1 activity and/or localization of Ipl1 substrates at the spindle, required for the disassembly, might be compromised in the absence of Sgo1. In support of this, the centromeric function of Aurora B, but not its localization, is regulated by shugoshin (Mei-S332) in *Drosophila* (Kerrebrock et al 1992, LeBlanc et al 1999, Resnick et al 2006) and in *Xenopus* (Sgo2) (Rivera et al 2012). Alternatively, CaSgo1 as adapter might recruit protein(s) at the microtubule plus ends residing at the spindle midzone and promote depolymerization of the microtubules. Similar to this, in vertebrates, shugoshin (Sgo2) has been shown to recruit a microtubule depolymerase, mitotic centromere-associated kinesin (MCAK) at the centromeres to regulate correction of erroneous KT-MT attachment by depolymerization of the plus ends (Huang et al 2007). Likewise, in budding yeast, proteins like Ase1 and Kip3 drive spindle disassembly through their spindle midzone localization (Juang et al 1997, Straight et al 1998, Gupta et al 2006, Varga et al 2006, Woodruff et al 2010). It would be interesting to examine if the homologous protein(s) in *C. albicans* act as effector(s) of spindle localized Sgo1. Currently, we are uncertain about how CaSgo1 might associate with the spindle. Since removal of the putative MT-binding domain did not perturb the deletion variant to go to the spindle (Fig. S2F, G), we believe that CaSgo1 might use either other domain for its association with the spindle or its association with other microtubule binding proteins and/or CPCs as in several organisms subcellular localization of shugoshin and CPCs are interdependent (reviewed in Marston 2015).

### Centromeric localization of shugoshin and function in bi-orientation and condensin recruitment

In various organisms, shugoshin has been observed first accumulating at the centromeres at prophase and then disappearing at anaphase during mitosis (reviewed in Marston 2015). For instance, Sgo1 remains undetected in α-factor arrested G1 cells in *S. cerevisiae*. ScSgo1 is expressed during early prophase stage, appears as a dot during metaphase but it becomes degraded during anaphase (Marston et al 2004, Nerusheva et al 2014). However, in *C. albicans,* we observed that CaSgo1 appears even at G1/S stage at the centromeres, as judged by the close proximity of Sgo1-2GFP to the spindle pole (Tub4), (Fig. 1B). Such interphase localization of shugoshin has been also observed for human Sgo1 where its centromeric localization is dependent on chromatin proteins including heterochromatin protein 1 (HP1, Yamagishi et al 2008, Kang et al 2011) which is, notably, irrespective of any KT-MT attachment. Consistent to this and converse to the notion that shugoshin accumulates at the MT-unattached tension-less centromeres, by ChIP assay we could detect Sgo1 at the centromeres that are attached and are under tension (Fig. 3A, Sgo1-TAP −Noc). In the absence of HP1 homolog in *C. albicans*, this pool of Sgo1 may recognize other centromeric chromatin mark and/or microtubule plus ends for the localization at the centromeres (Fig. 3C, TENSION on, SAC off). On the other hand, as commonly observed in other organisms, we could also detect an enrichment of Sgo1 at the centromeres when they are not under tension (Fig. 3A, Sgo1-TAP + Noc). This pool of Sgo1 presumably follows Bub1-H2A pathway (reviewed in Marston 2015) for its enriched localization at the unattached centromeres (Fig. 3C, TENSION off, SAC on). In other organisms, shugoshin localized at the unattached centromeres acts as an adapter to sense lack of tension and promotes sister chromatid bi-orientation through recruitment of CPC (reviewed in Marston 2015). These functions of shugoshin appear to be conserved in *C. albicans* (Fig. 3A, 6 and 7D). Additionally, in budding yeast, ScSgo1 promotes bi-orientation and error-correction in KT-MT attachment by recruiting condensin and maintaining Ipl1 at the centromeres (Verzijlbergen et al 2014, Peplowska et al 2014). Remarkably, our finding that CaSgo1 also facilitates condensin localization and Ipl1 maintenance at the centromeres (Fig. 7C, D) suggests that a similar theme might be adopted to geometrically constrain ‘back-to-back’ orientation of the sister kinetochores in *C. albicans*. Compared to an organism with point centromere, in an organism with epigenetically determined regional centromeres (Sanyal et al 2004, Baum et al 2006), the pericentromeric chromatin architecture is expected to play more significant role to drive faithful chromosome segregation. Therefore, it would be interesting to investigate if shugoshin, through condensin targeting, might influence the architecture of the pericentromeres in *C. albicans* having regional centromeres. Such a possibility also stems from the finding that an alteration in pericentric chromatin organization has been observed in budding yeast lacking ScSgo1 (Haase et al 2012).

### Shugoshin as an activator of SAC

Although the function of shugoshin as a sensor for the lack of tension appears to be conserved, the interaction of shugoshin with SAC for the purpose of onward signaling of the response varies in a species-specific manner. In human, mouse and frog, shugoshin (Sgo2 or Sgo) has been shown to physically interact with SAC protein, Mad2 in meiosis but not in mitosis (Orth et al 2011, Rattani et al 2013). Given this physical interaction and the evidence of forming Sgo-closed-Mad2 complex, it is intriguing to examine whether shugoshin acts upstream or downstream of Mad2. This appears to vary amongst the species. From the finding that loss of Sgo1 in HeLA cells caused persistent association of Mad2 and BubR1 at the centromeres with increased mitotic arrest (Salic et al 2004, McGuinness et al 2005, Tang et al 2004), it appears that in mitosis shugoshin acts downstream of Mad2, akin to Cdc20. This appears to be also true in meiosis as in human, depletion of Mad2 caused reduced Sgo2 localization at the centromeres and a mutant variant of Sgo2 unable to interact with Mad2, was observed less focused at the centromeres (Orth et al 2011). On the other hand, shugoshin, like Mad1, acting as upstream positive regulator of Mad2 (or SAC) has also been demonstrated in some instances. In Sgo2-null mice, several spermatocytes escape meiosis II arrest and produces aneuploid spermatozoa which is proposed due to lack of activation of SAC (Llano et al 2008). Recently, in *Xenopus,* the Sgo1-Mad2 and in humans the Sgo2-Mad2 interactions have been shown to inhibit separase in a securin-independent manner (Hellmuth et al 2020).

In budding yeast, revelation that ScSgo1 functions as tension sensing component of SAC, it appears that in this organism ScSgo1 also acts upstream of SAC. Consistent to this, removal of Mad2 does not reduce the level of ScSgo1 at the centromeres (Nerusheva et al 2014). However, ScSgo1 is dispensable for SAC mediated metaphase arrest upon microtubule removal (Indjeian et al 2005, Nerusheva et al 2014) indicating that ScSgo1 does not directly activate SAC by recruiting and or maintaining it at the centromeres. Since there is no abrogation of metaphase arrest, ScSgo1 is not recognized as a SAC component but rather has been shown later to be a part of SAC silencing network (Jin et al 2013). In contrast, in *C. albicans* we show that, the SAC-mediated metaphase arrest was compromised in CaSgo1-null cells harboring defects in KT-MT attachments (Fig. 4A, Suppl. Fig. S5B). Possibly, the observed drop in cell viability of the Ca*sgo1* mutant following exposure to anti-microtubule drug (Fig. 4D, Suppl. Fig. S5A) might be, akin to *S. cerevisiae*, due to failure in de novo sister chromatid bi-orientation during spindle recovery from the drug. Nevertheless, the generation of multibudded cells in presence of the drug (Suppl. Fig. S5B), argues for attenuation of SAC and bypass of the mitotic arrest in absence of Sgo1 in *C. albicans*. These results indicate that in *C. albicans* Sgo1 plays a role for the activation of SAC at the centromeres. Consistent to this we observed that the maintenance of SAC, at least Mad2 and Bub1, is compromised in Sgo1-null cells (Fig. 5). Interestingly, we also observed a reduction in Sgo1-GFP intensity at the centromeres in Mad2-null cells in absence of tension (Fig. 3B), indicating that localizations of Sgo1 and Mad2 are interdependent in *C. albicans*. This implies that in this organism, in contrast to other model yeasts, Sgo1 plays a greater role in SAC related functions. On the other hand, a synergistic effect between *sgo1* and *mad2* was observed under anti-microtubule drug challenge (Suppl. Fig. S6B, C) suggesting, that CaSgo1 might have additional SAC-independent function(s) to inhibit anaphase promoting complex (APC) in response to the drug to cause cell cycle delay. Such a SAC independent function of delaying cell cycle by ScSgo1 has been reported in budding yeast in case of lack of tension. In that case it is believed that ScSgo1 recruited PP2A antagonizes polo kinase (Cdc5) mediated phosphorylation of cohesin (Scc1), that blocks its cleavage from separase (Yaakov et al 2012) or PP2A might directly resist separase activity (Clift et al 2009). These results altogether indicate that in *C. albicans* shugoshin may function as a component of SAC and/or activate the SAC when KT-MT attachments are compromised.

Overall, in this study, we characterize shugoshin for the first time in *C. albicans*. Consistent to the conserved functions of shugoshin across eukaryotes, we observed several mitotic functions which are similar to other organisms (Fig. 8). Notably, we report here two important adapter functions along the cell cycle that have not been yet reported in any other organisms including fungi (Fig. 8). First, shugoshin in *C. albicans* may play additional roles to regulate genome segregation under stress by maintaining and activating SAC at the centromeres. It may not be surprising if an induced attenuation of CaSgo1 function under stress conditions occurs to license certain aneuploidy enforcing stress tolerance. Second, with ever increasing range of adapter functions of shugoshin, our finding that CaSgo1 associates with the microtubules in vivo uncovers a novel function of this protein at the spindle besides what has been observed till date at the chromosomes and at the centrosomes. Taken together, this study reports a conserved principle of shugoshin function across organisms harboring genetically and epigenetically defined centromeres, with evidence of novel functions in *C. albicans.*

**Figure 8.**
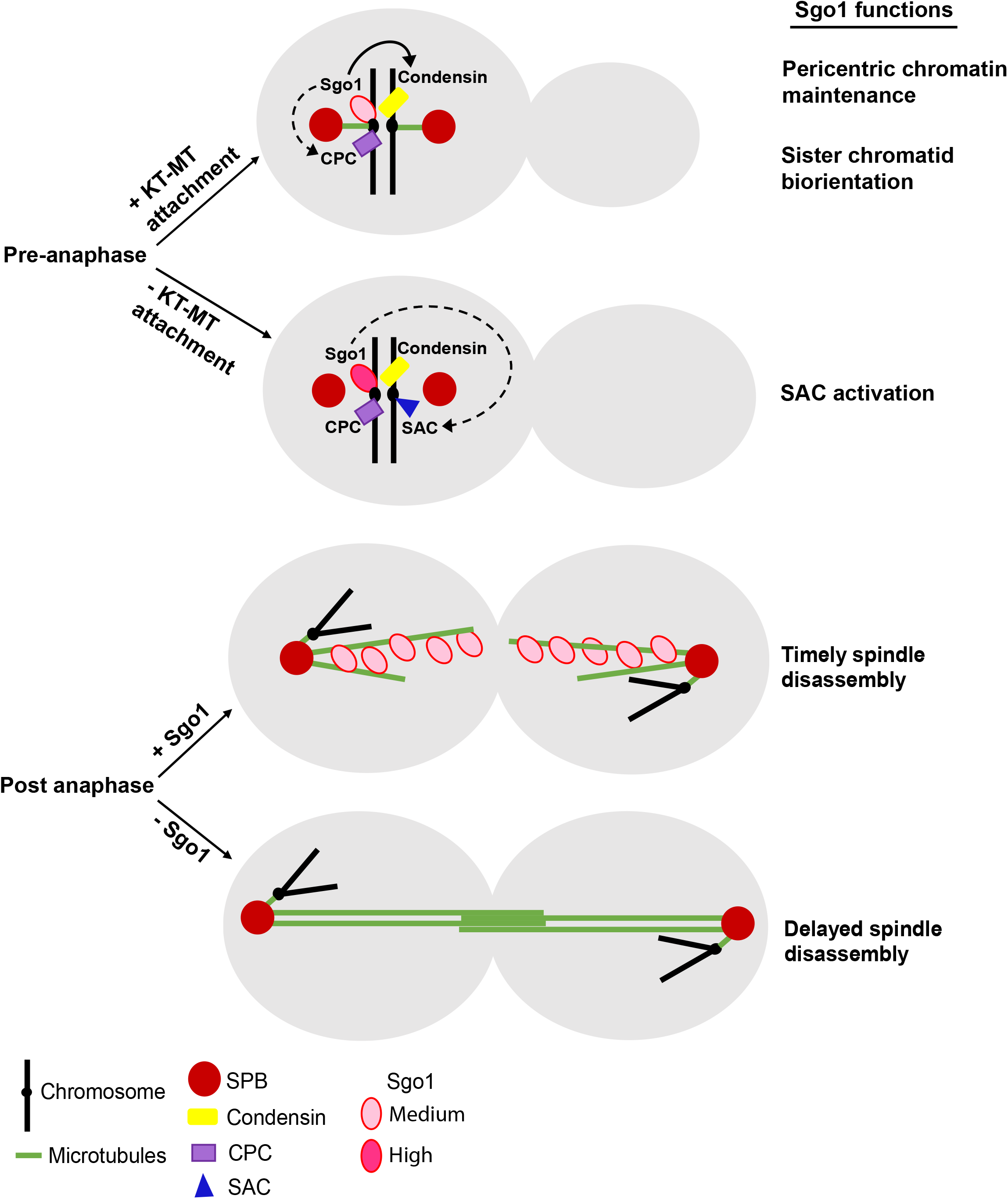
Schematic depicting Sgo1 functions in *C. albicans*. During pre-anaphase, under normal condition (+KT-MT attachment), Sgo1 maintains pericentric chromatin by condensin recruitment (solid arrow) and promotes chromosome biorientation through correction of erroneous kinetochore-microtubule attachment by maintaining (dotted arrow) CPC, at the centromeres/pericentromeres. Upon loss of tension between the sister kinetochores (- KT-MT attachment), Sgo1 functions as a SAC component by maintaining the SAC proteins at the centromeres to facilitate cell cycle arrest. During post anaphase, Sgo1 exhibits a microtubule-dependent spindle localization and ensures a timely spindle disassembly.

Given, both mutant and overexpression alleles of shugoshin have been implicated in human diseases including various cancers and alteration in heart and gut rhythm (Iwaizumi et al 2009, Liu et al 2012, Yamada et al 2012, Matsuura et al 2013, Yang et al 2013, Chetaille et al 2014), it is imperative to reveal all the functions of shugoshin by studying various organisms. On the other hand, our findings on *C. albicans* and use of Sgo1 as a potential therapeutic target for hepatocellular carcinoma (Wang et al 2015) may pave the way for exploring CaSgo1 as a potential anti-Candida drug target in the near future.

## Materials and methods

### Strains, plasmids and primers

All the *C. albicans* strains, bacterial plasmids and primers used in this study are mentioned in Table S1, Table S2 and Table S3, respectively in the supplementary material.

### Construction of heterozygous and conditional mutant strains

For generating *sgo1* heterozygous mutant SGY8001, one copy of *SGO1* in strain SN148 was replaced with *SAT1* flipper cassette using KpnI/SacI fragment from the plasmid pSFS2a-*SGO1*-USDS which was constructed by cloning upstream and downstream sequences of *SGO1*, flanking the *SAT1* flipper cassette in pSFS2a. Following deletion of *SGO1*, *NAT* marker from the cassette was rescued.

For generating conditional mutant, the native promoter of the second copy of *SGO1* in strain SGY8001 was shuffled with *MET3* promoter by linearizing (using BstBI site in *SGO1*) and integrating pCaDIS-*SGO1*, which was constructed by cloning upstream sequence (1283 bp) of *SGO1* as BamH1-Pst1 fragment in-frame with *MET3* promoter sequence in pCaDIS.

### Construction of homozygous mutant and rescue strains

For construction of *sgo1* homozygous null mutant, the second copy of *SGO1* in strain SGY8001 was replaced with SAT1 flipper as mentioned above, generating strain SGY8035 from which the *NAT* marker was rescued, generating strain SGY8037. Strains SN148, SGY8001, SGY8035 and SGY8037 were confirmed for *SGO1* deletions by Southern hybridization. URA3 was integrated in these strains using the plasmid CIp10.

For constructing *SGO1* rescue strain, full length *SGO1* including upstream and downstream sequences was cloned in CIp10 to generate plasmid CIp10-*SGO1-URA3* which was then linearized at RP10 locus by Stu1 and integrated in strain SGY8037 to generate strain SGY8116.

For constructing *sgo1* homozygous null mutant in YJB13024 strain, above strategy was used.

### Construction of epitope tagged strains

For *SGO1-2GFP* tagging, we first constructed plasmid pBS-*SGO1-GFP-HIS1* by cloning *SGO1* in pBS-*GFP-HIS1* plasmid. We next cloned *GFP*-linker in the above plasmid to generate pBS-*SGO1-2GFP-HIS1*. This was then linearized within *SGO1* by partial restriction digestion using BstB1 and was integrated into strains SN148 and J110 to generate strains SGY8156 and SGY8243, respectively.

For constructing *SGO1MTΔ-GFP* allele, we first amplified *SGO1MTΔ* sequence containing 300 bp deletion (from +772 nt to +1072 nt) within *SGO1* ORF, by overlap extension PCR using primers SA33, SA82 and SA85, SA34. We then cloned *SGO1MTΔ* sequence in pBS-*GFP-HIS1* plasmid to generate pBS-*SGO1MTΔ-GFP-HIS1*. This was then linearized within *SGO1* restriction digestion using AatII and was integrated into strain SGY8001 to construct SGY8331 strain (Table S1).

For *SGO1-TAP* tagging, a Candida optimized tandem affinity purification (TAP) tag cassette was amplified from plasmids pFA-*TAP-ARG4* and pFA-*TAP-HIS1* using primers SA6 and SA7 and strains SN148 and J110 were transformed with this cassette to replace one copy of *SGO1* with *SGO1-TAP*, generating strains SGY8038 and SGY8130, respectively. For tagging the second copy of *SGO1* in strain SGY8038, same strategy was followed using plasmid pFA-*TAP-HIS1*.

For *MAD2-3HA* and *Smc4-3HA* tagging, above strategy was followed using plasmid pFA-*HA-ARG4* and primers SA69, SA70 and SA65, SA66 respectively to generate the indicated strains (Table S1).

For *TUB1-GFP* tagging, one copy of *TUB1* was tagged with GFP by linearizing and integrating plasmid pBS-*TUB1-GFP-HIS1*, which was constructed by cloning *TUB1* fragment (SA29, SA30) in pBS-*GFP-HIS1* (SacII, SpeI) plasmid, to generate the indicated strains (Table S1).

For *IPL1-2GFP* and *TUB1-RFP*, one copy of *IPL1* and *TUB1* was tagged with 2GFP and RFP respectively, by linearizing (XbaI) and integrating plasmids pBS-*IPL1-2GFP-HIS1* and pBS-*TUB1-RFP*, respectively to generate the indicated strains (Table S1).

For *TUB4-MCHERRY* tagging, one copy of *TUB4* was tagged with *MCHERRY* by linearizing (BstEII) and integrating plasmid CIp10-*TUB4-MCHERRY-URA3* or plasmid CIp10-*TUB4-MCHERRY-LEU2* to construct the indicated strains (Table S1)

### Media and growth conditions

All cultures were grown in liquid media (YEPD) or solid media (YEPD agar) supplemented with 100 μg/ml uridine, at 30°C shaking (200 rpm) or static respectively, unless stated otherwise.

For conditional repression of *SGO1*, all strains were grown overnight, reinoculated at 0.2 OD_600_ in YEPD+100 μg/ml uridine and harvested at 0.8 OD_600_ serially diluted and spotted on synthetic complete media (CM) plates which were supplemented without or with the indicated concentrations of cysteine and methionine.

For conditional repression of *DAM1*, all strains were treated as described earlier (Thakur et al 2011) for the indicated time points in Dam1 permissive (-CM) and repressive (+CM) media respectively.

### Drug sensitivity and viability assay

For nocodazole sensitivity assay, all strains were treated without (−NOC) or with (+NOC) 20 μg/ml Nocodazole (Sigma, M1404) as described earlier (Bai et al 2002, Thakur and Sanyal, 2011) for the indicated time points, harvested, washed with sterile water and spotted on drug-free synthetic complete media. For viability assay (quantification) same procedure was followed except, that the cells were plated on drug-free synthetic complete media.

For hydroxyurea (HU) (Sigma, H8627) sensitivity assay, all strains were grown overnight, reinoculated at 0.2 OD_600_ in YEPD+100 μg/ml uridine grown for one generation (0.4 OD_600_) treated with 20 mM HU for 1 hr (Bai et al 2002), before harvesting, washing and spotting on drug-free YEPD agar plates.

### Bi-orientation assay

Chromosome bi-orientation assay was performed in *C. albicans* as described previously for *S. cerevisiae* (Fernius and Hardwick, 2007) with some modifications. Briefly, midlog grown cultures of cells were treated with 20 μg/ml nocodazole for 2 h. The cells were harvested, washed, and were allowed to grow in drug free medium. The cells were harvested after 15 and 30 min to analyze bi-orientation of chromosomes and sister chromatid disjunction, respectively. To verify the condition of the spindle before and after nocodazole treatment and during recovery process, tubulin was visualized by immunofluorescence assay (see below) as described earlier (Mehta et al 2014).

### Live cell DAPI staining for sister chromatid cohesion (SCC) assay

DAPI (Invitrogen, D1306) was added at the final concentration of 7 μg/ml to the early log culture and the cells were allowed to grow for one more generation. Subsequently, the cells were grown for 2 h in the absence (−NOC) or presence (+NOC) of 20 μg/ml nocodazole. Cells were then harvested, washed and imaged.

### Live cell microscopy and image processing

All live cell microscopy images, except for bi-orientation assay, were captured using confocal laser scanning microscope LSM 780. Images were processed using Zeiss Zen 2012 software. All intensity measurements of fluorescence signals were done using ImageJ software and intensity values (a.u) reported after correction with the background signal. Imaging for bi-orientation assay was done using AxioObserver Z1 Zeiss inverted microscope and processed using AxioVs40 V 4.8.2.0 software.

### Indirect Immunofluorescence

This assay was performed as described previously (Mehta et al 2014) with some modifications. For spheroplasting, cells were treated with zymolyase 20T (MP Biomedicals, 320921) in the presence of 25 mM β-mercaptoethanol for 20 min at 30°C. For immuno-staining, rat anti-tubulin antibodies (1:500 dilution; Serotech YOL1/34, MCA78G) followed by TRITC goat anti-rat antibodies (1:500 dilution; Jackson, 115-485-166) were used. DAPI (1 μg/ml) was used for nuclear staining.

### Chromatin spread

Chromatin spread assay was performed as described previously (Mehta et al 2014) with some modifications. For fixing spheroplasted cells on acid washed glass slides, instead of using 4% paraformaldehyde solution, 3% paraformaldehyde solution was used to maximize the spreading efficiency. Mouse anti-GFP (1:200 dilution; Roche, 11814460001) was used for staining Cse4-GFP and rat anti-HA (1:200 dilution: Roche, 11867423001) for Mad2-3HA staining as primary antibodies. Secondary antibodies used were Alexa-fluor 488 goat anti-mouse (1:500 dilution; Jackson, 115-545-166) and TRITC goat anti-rat (1:500 dilution; Jackson, 112-025-167) respectively; and DAPI for nuclear staining.

### Flow cytometry

Flow cytometry analysis was performed as previously described (Agarwal et al 2015) in the strains at indicated time points of without (−CM) or with (+CM) Dam1 depletion using propidium iodide for nuclear staining.

### Western blotting

For protein extraction, all strains were grown overnight, reinoculated in at 0.2 OD_600_ in 50 ml of YEPD+100 μg/ml uridine media, grown till 0.8-1 OD_600_ and harvested, washed once with distilled water, and dried pellets were kept in boiling water bath for 10 min. Pellets were then resuspended in 80 μl of ESB buffer (1.5 mM TRIS-HCl, pH 6.8, 2 M DTT, 20% SDS, 0.3% bromophenol blue and 50% glycerol) containing 1X protease inhibitor cocktail (Roche, 11873580001), to which 100 μl acid-washed glass beads (0.5 mm) were added. This mix was vortexed 5 times, with 1 min on vortexer and 1 min on ice. More 60 μl of ESB buffer was added followed by 1 min vortexing. Beads and cell debris were separated from supernatant by centrifuging at 3000 rpm for 1 min. Lysate was cleared by centrifugation at 10000 rpm for 30 s, supernatant collected, boiled in the presence of SDS loading dye and subjected to electrophoresis on 8-10% SDS PAGE. Post this, protein samples were transferred to nitrocellulose membrane (Pall Lifesciences, BSP0161), membrane blocking was done with 5% skimmed milk in TBST solution and subjected to epitope-specific primary antibodies and HRP-conjugated secondary antibodies. Development of blots were done using ECL substrate (Biorad, 1705060).

### Southern hybridization

Southern blotting was performed as described previously (Varshney et al 2019). Primers P24, P25 were used for preparation of amplicon which was used for probe preparation. Restriction enzyme EcoRV was used for the digesting the genome of indicated strains.

### Chromatin immunoprecipitation and qPCR

Chromatin immunoprecipitation assay was performed as described elsewhere (Prasad et al 2019, Mehta et al 2014, Burrack et al 2013) with modifications. Formaldehyde cross-linking of Sgo1-TAP and Smc4-3HA proteins were done for 20 min and 1 h 15 min respectively at RT, followed by overnight incubation at 4°C. For pulldown of DNA-protein complex, protein-A Sepharose (G.E healthcare, 17078001) beads were precoated with anti-protein A antibodies (Sigma, P3775) and anti-HA antibodies (Roche, 11867423001) by adding 5 μg of antibodies (+Ab) or without antibodies (-Ab) in 50 μl of beads resuspended in 250 μl of 1X PBS containing 5 mg/ml BSA, and this mix was added to the lysate. Enrichment/Input values obtained by qPCR as described previously (Mehta et al 2014). All the enrichment/input values obtained in -Ab samples were subtracted from +Ab samples to eliminate any non-specific enrichment of DNA fragments by beads. Error bar of each ChIP sample indicates standard error calculated from three independent biological replicates. Statistical significance is denoted by calculating *p* value using two-tailed Student’s *t*-test.

### Statistical analyses

Error bar of each ‘type’ indicates standard error calculated from three technical replicates using at least two biological replicates. Statistical significance is denoted by calculating *p* value using two-tailed Student’s *t*-test.

## Abbreviations

KT-MT: kinetochore microtubule
SAC: Spindle assembly checkpoint
SPB: spindle pole body
CPC: chromosomal passenger complex

## Author contributions

AS performed all the experiments, analysis of the data and writing of the manuscript. SS constructed Bub1-GFP strain. KS and SKG contributed in analysis of the data and writing/editing of the manuscript. KS and SKG gathered funding. SKG supervised the project.

## Acknowledgements

We thank J. Berman, D. Kirkpatrick for providing plasmids and strains of *C. albicans.* We thank members of SKG Lab and KS Lab for helpful discussions about the experiments. This project was supported by a grant from the Department of Biotechnology (BT/PR13909/BRB/10/1432/2015) to SKG and KS. KS is a Tata Innovation Fellow. AS is supported by MHRD (144300004) and IIT Bombay for her fellowship. S.S. thanks Council for Scientific and Industrial Research (09/733(0192)/2014-EMR-I) and JNCASR for his fellowship. We gratefully acknowledge FACS facility and Confocal Laser Scanning microscope of the Central Instrumentation Facility of IIT Bombay.

## Conflict of interest statement

The authors have declared that they have no conflicts of interest.

## Supplementary Figure Legends

**Figure S1. Conserved sequence features of *SGO1* in *C. albicans* and phylogenetic analysis of shugoshin in species from the Ascomycota phylum.**

(A) Amino-acid sequence alignment of shugoshin from different organisms. Conserved motifs and/or amino-acids are highlighted using BLASTP, CLUSTALW and GBLOCKS sequence analysis tools. (B) Schematic of conserved structural motifs found in shugoshin proteins. Dm: *Drosophila melanogaster*, Hs: *Homo sapiens*, Mm: *Mus musculus*, Xl: *Xenopus laevis*, Sp: *Schizosaccharomyces pombe,* Sc: *Saccharomyces cerevisiae*, Ca: *Candida albicans*. (C) Out of ~30000 ascomycetes species, in 437 species shugoshin has been annotated in NCBI. Amino acid sequences of shugoshins from these species were retrieved from NCBI. Each leaf corresponds to an accession ID. Tree was constructed by alignment of the sequences using Clustal Omega MSA and iTOL (Interactive Tree of Life). 93 species (majorly, but not exclusively, from the Saccharomycotina sub-phylum) sharing an internal node, from the indicated sub-phylum, have been highlighted to show the placement of CaSgo1 with respect to the other ascomycetes in the group. Black arrowhead shows an internal node shared by CaSgo1 and ScSgo1 suggestive of a recent common ancestor. (D) Comparison of the sequences of CaSgo1, ScSgo1 and SpSgo1 highlights (arrowhead) the common ancestor of CaSgo1 and ScSgo1 depicted by the internal node. In (C) and (D), black lines and red line (used to highlight CaSgo1) indicate sequences annotated as ‘Sgo1’ or ‘shugoshin’ in the database, whereas blue lines indicate the paralog ‘Sgo2’.

**Figure S2. Sgo1 is non-essential for mitotic growth under unperturbed conditions, Sgo1 fusion alleles are functional, and the MT binding motif of Sgo1 is dispensable for its spindle localization.**

(A) The cell viability of the strain SGY8005 (*sgo1Δ/P*_*MET3*_*SGO1*) as compared to the wild type strain SN148 (*SGO1/SGO1*) in absence (-CM) or presence (+CM) of increasing concentrations of methionine and cysteine. Strain SGC65 (*sth1Δ/P*_*MET*_*STH1*) containing conditionally repressible essential gene *STH1* was used as a control. (B) *Right*, Southern analysis confirming sequential deletion of both the copies of *SGO1* in the strain SGY8037 (*sgo1Δ/Δ*). *Left*, schematic depicting position of the probe binding and different restriction fragment lengths. (C) *Left*, growth curve of the wild type and *sgo1Δ/Δ* mutant cells carried out at 30°C. *Right*, percentage of the cells depicting different cell cycle stages in the exponentially growing cultures of the wild type and *sgo1Δ/Δ* mutant. The number of cells analysed, denoted here and in subsequent experiments as *n*, were at least 100 (*n* ≥ 100); ‘abnormal’ population was compared between the strains to calculate *P* values. (D) Wild type strain SN148 (*SGO1/SGO1*), SGY8095 (*SGO1-TAP/SGO1-TAP*) wherein both the copies of *SGO1* were fused with TAP tag, and SGY8037 (*sgo1Δ/Δ*) were mock treated (−NOC) or treated with 20 μg/ml nocodazole (+NOC) and spotted on drug-free medium plates. The plates were incubated for 36 h at 30°C before they were photographed. (E) Same as in ‘D’, except using the strains SGY8264 (*SGO1/SGO1::TUB4/TUB4-MCHERRY*), Sgo1-2GFP strain SGY8303 (*sgo1Δ/SGO1-2GFP::TUB4/TUB4-MCHERRY*) and *sgo1Δ/Δ* mutant SGY8305 (*sgo1Δ/Δ::TUB4/TUB4-MCHERRY*). (F) Schematic representation of GFP fused full length Sgo1 (above) and deletion construct of Sgo1 lacking amino acid residues 258 to 358 designated as microtubule (MT) binding motif (below). (G) Fluorescence images of live cells using strain SGY8330 (*sgo1Δ/SGO1-GFP::TUB1/TUB1-RFP)* and SGY8331 (*sgo1Δ/SGO1 MTΔ-GFP::TUB1/TUB1-RFP)* depicting localization of Sgo1-GFP or Sgo1 MTΔ-GFP with respect to MTOC (type I) and spindles (type II) marked by Tub1-RFP. *n* ≥ 140; type II population was compared between the strains to calculate *P* values scale bar ~ 5 μm. Histogram plot depicts distribution of type I and II categories among the metaphase and anaphase cell population of the indicated strains. Error bars, in this and subsequent experiments, denote standard error calculated from standard deviation obtained from three experimental replicates; asterisk (*) and (**ns) denote statistically significant (*P* < 0.05) and non-significant (*P* > 0.05) values respectively, calculated using two-tailed Student’s *t*-test.

**Figure S3. *sgo1Δ/Δ* cells do not show any gross abnormality in spindle morphology.**

(A) Live cell imaging of the microtubule spindle (Tub1-GFP) using wild type SGY8171 (*SGO1/SGO1::TUB1/TUB1-GFP::TUB4/TUB4-MCHERRY*) and mutant SGY8172 (*sgo1Δ/Δ::TUB1/TUB1-GFP::TUB4/TUB4-MCHERRY*) strains. The histogram plot displays distribution of cells exhibiting dot-like (type I), short (type II) and anaphase (type III) spindles among the cell population; *n* ≥ 101. Scale bar ~5μm (B) Box plot depicting spindle lengths in metaphase cells of the strains used in (A); *n* = 53. (C) Box plot depicting background corrected fluorescence intensities of Ipl1-2GFP at the anaphase spindle (SPB-SPB distance >4 μm) in strains SGY8284 (*SGO1/SGO1::IPL1/IPL1-2GFP::TUB4/TUB4-MCHERRY*) and SGY8290 (*sgo1Δ/Δ::IPL1/IPL1-2GFP::TUB4/TUB4-MCHERRY)*; *n* = 20. **ns = statistically non-significant (*P* > 0.05).

**Figure S4. Expression of Sgo1 protein in cycling *C. albicans* cells.**

Whole cell extracts of SN148 (*SGO1/SGO1*) and SGY8038 (*SGO1/SGO1-TAP*) strains were separated on SDS PAGE. An estimated 80 kDa band of Sgo1-TAP was detected by anti-Protein A antibodies on western blot. Tubulin was used as a loading control, asterisk (*) indicates nonspecific bands developed by anti-α-tubulin antibodies.

**Figure S5. Sgo1 is required for SAC surveillance and is dispensable for DNA damage response in *C. albicans*.**

(A) Nocodazole sensitivity assay of SGY8114 (*sgo1Δ/Δ*) and SGY8112 (*sgo1Δ/SGO1)* as compared to the wild type SGY8110 (*SGO1/SGO1*) strains. J110 (*mad2Δ/Δ*) and the *SGO1-* add back rescue strain SGY8116 (*sgo1Δ/Δ:SGO1)* were used as controls. All the strains were treated with 20 μg/ml nocodazole or DMSO for the indicated time, before the cells were harvested, washed and serially diluted cultures were spotted on SD agar plates without nocodazole. (B) Strains same as (A) were analysed for nuclear segregation following treatment with nocodazole for indicated time points. SGY8114 (*sgo1Δ/Δ*) and J110 (*mad2Δ/Δ*) mutants formed multi-budded (arrowheads) cells from 4 h onwards as compared to the wild type SGY8110 (*SGO1/SGO1*) cells which became arrested in 2 h and started elongating (arrows) in subsequent time points. Scale bar ~5 μm. (C) Hydroxyurea (HU) sensitivity assay of the mutant SGY8114 (*sgo1Δ/Δ*) and *SGO1-*add back rescue strain SGY8116 (*sgo1Δ/Δ::SGO1*) compared to the wild type SGY8112 (*SGO1/SGO1*). Strains DKCa596 (*mec1Δ/Δ*) and J110 (*mad2Δ/Δ*) were used as controls. The cells were pretreated with 20 mM HU for 60 min before they were washed and spotted on drug free SD media plates.

**Figure S6. Synergistic effect between *sgo1Δ/Δ* and *mad2Δ/Δ*.**

(A) Western blot analyses depicting expression of Mad2-3HA in the wild type SGY8301 (*CSE4/CSE4-GFP-CSE4::MAD2/MAD2-3HA*) and *sgo1Δ/Δ* mutant strain SGY8302 (*sgo1Δ/Δ*::*CSE4/CSE4-GFP-CSE4::MAD2/MAD2-3HA*) wherein one copy of *MAD2* was tagged C-terminally with 3HA. (B) Nuclear segregation and budding index analyses of the cells following nocodazole treatment for the indicated durations in the double mutant strain SGY8322 (*sgo1Δ/Δ::mad2Δ/Δ*) as compared to the corresponding single mutants SGY8114 (*sgo1Δ/Δ)*, SGY8318 (*mad2Δ/Δ*) and wild type SGY8110 (*SGO1/SGO1*). Arrows and arrowheads indicate large budded arrested cells and arrest bypassed cells, respectively; *n* for 0 h ≥ 38, *n* for 2 h onwards ≥ 154, ‘multibudded’ population was compared between the strains to calculate *P* values. Scale bar ~5 μm (C) Viability assay of the strains used in (B) for the indicated duration of nocodazole treatment.

**Figure S7. *sgo1Δ/Δ* cells show increased chromosome non-disjunction during spindle recovery.**

SGY8154 (wild type) and SGY8165 (*sgo1Δ/Δ*) cells were treated with 20 μg/ml nocodazole for 90 min and were released into drug-free media for 30 min before they were imaged to assess sister chromatids disjunction. (A) Immunofluorescence staining of tubulin in the indicated strains and conditions to visualize successful recovery of tubulin after 15 min into drug free medium. (B) Cells recovered for 30 min were harvested to score sister chromatids disjunction. Representative images of the cells showing sister chromatids disjunction (type I), scored as one GFP dot each in mother and daughter cells and non-disjunction (type II), scored as one or two GFP dots in either mother or daughter cells. Scale bar ~5 μm. Histogram plot depicts the distribution of type I and II categories among the cell population; *n* ≥ 185.

**Figure S8. Loss of Sgo1 does not affect the initial centromeric localization of Ipl1.**

(A) Western blot analysis showing the expression of a condensin subunit Smc4-3HA in wild type SGY8238 (*SMC4/SMC4-3HA*) and *sgo1Δ/Δ* mutant SGY8227 (*sgo1Δ/Δ::SMC4/SMC4-3HA*). No tag strain SN148 (*SMC4/SMC4*) was used as control. (B) Live cell imaging of wild type SGY8284 (*IPL1/IPL1-2GFP::TUB4/TUB4-MCHERRY*) and *sgo1Δ/Δ* mutant SGY8290 (*sgo1Δ/Δ::IPL1/IPL1-2GFP::TUB4/TUB4-MCHERRY)* cells at start of the cell cycle before separation of SPBs (Tub4). Localization of Ipl1 as tight knit dot (localized) next to SPBs was considered as centromeric localization. Percentage of cells from the indicated strains with localized or no signal of Ipl1 are shown on right; *n* ≥ 100. Scale bar ~5 μm.

